# TMEDs Mediate Versatile Cargo Transport in Vesicle-dependent Unconventional Secretion

**DOI:** 10.1101/2025.05.04.652080

**Authors:** Jianfei Zheng, Haodong Wang, Yuxin Sun, Pei Chang, Xing Deng, Lijingyao Zhang, Lin Zhu, Kangling Zhu, Haiteng Deng, Min Zhang, Liang Ge

**Affiliations:** State Key Laboratory of Membrane Biology, Tsinghua-Peking Center for Life Sciences, Beijing Frontier Research Center for Biological Structure, School of Life Sciences, Tsinghua University, Beijing, 100084, China; School of Pharmaceutical Sciences, Tsinghua University, Beijing, 100084, China; Academy for Advanced Interdisciplinary Studies, Beijing, Peking University, 100871, China.; MOE Key Laboratory of Bioinformatics, Beijing, China

## Abstract

Unconventional protein secretion (UcPS) exports diverse signal peptide-lacking cargoes, yet its cargo selectivity remains poorly understood. Here, we identify TMED proteins as key regulators of vesicle-dependent UcPS, mediating selective cargo release via translocation into secretory carriers. TMED proteins act as translocators, facilitating cargo passage across lipid bilayers with assistance from HSP90 chaperones and partial cargo unfolding. Selectivity arises during translocation, where TMED cytoplasmic tails bind specific cargoes. The ER-Golgi intermediate compartment (ERGIC) is essential for TMED-mediated translocation and release. TMED homo-oligomerization enhances translocation, while hetero-tetramerization inhibits it. ERGIC localization promotes homo-oligomerization, which is further stabilized by cargo binding, forming a feed-forward mechanism to enhance translocation. These findings establish TMED proteins as central regulators of cargo diversity in UcPS, with their oligomerization and subcellular localization modulating translocation efficiency.

## Introduction

Protein secretion constitutes a fundamental biological mechanism facilitating intercellular communication and signal transduction. In eukaryotic cells, numerous secretory proteins feature a signal peptide that enables their recognition by the signal recognition particle (SRP), followed by translocation into the endoplasmic reticulum (ER) via the SEC61 translocon (Rapoport et al., 2017; Shan and Walter, 2005; Voorhees and Hegde, 2016), and subsequently, proceed through ER-Golgi trafficking (Pantazopoulou and Glick, 2019; Zanetti et al., 2011) , the process of which is termed conventional secretion. However, recent research has unveiled unconventional protein secretion (UcPS) pathways, in which cytosolic proteins lacking a signal peptide are secreted independently of the ER-Golgi route (Rabouille, 2017; Rabouille et al., 2012; Zheng and Ge, 2022). These UcPS pathways play a role in a variety of biological processes encompassing inflammation, development, and metabolism (Claude-Taupin et al., 2017; Pallotta and Nickel, 2020; Prentice et al., 2021; Villeneuve et al., 2018). Disruptions in UcPS or the release of mutant cargoes have been linked to human diseases, including neurodegenerative disorders and cancer (Ejlerskov et al., 2013; Lock et al., 2014).

An essential inquiry pertains to the differential release of various UcPS cargoes under specific conditions. While various UcPS pathways have been identified, including both vesicle-independent and vesicle-dependent routes (Rabouille, 2017; Sparn et al., 2022; Zheng and Ge, 2022), it remains a significant challenge to elucidate the precise mechanisms governing the distinct control of diverse cargoes by these pathways. In the context of vesicle-mediated UcPS, a pivotal stage involves the regulation of cargo entry into the secretory vesicle, as this stage determines the transition of a cytosolic protein into a secretory protein. However, the mechanisms underlying how a UcPS cargo, lacking a signal peptide, translocates into a secretory vesicle and how the translocation system accommodates cargo diversity remain largely uncharted.

Previously, we identified a translocation event that regulates the entry of several UcPS cargoes into the ER-Golgi intermediate compartment (ERGIC) as an essential step for vesicle-mediated UcPS which likely then traffick vesicles that partially overlap with pathways of secretory autophagy and multi-vesicular body trafficking (Dupont et al., 2011; Liu et al., 2020; Zhang et al., 2015; Zhang et al., 2020). The transmembrane emp24 domain-containing protein 10 (TMED10/TMP21) was implicated as a potential protein channel (or translocator) accounting for the translocation of UcPS cargoes into the ERGIC and therefore the pathway was tentatively termed as the TMED10-channeled UcPS (THU) (Liu et al., 2020; Zhang et al., 2015; Zhang et al., 2020). Multiple physiological roles of THU have recently been identified, including the regulation of intestinal epithelium differentiation (Wang et al., 2024), neuronal homeostasis (Jiao et al., 2024), cancer immunity (Zhang et al., 2024), and inflammation associated with coronavirus infection (Liu et al., 2024). Notably, THU mediates the release of various types of UcPS cargoes under distinct physiological and pathological conditions. However, with the growing list of identified UcPS cargoes, it remains uncertain how THU, predominantly driven by the relatively simple structure of TMED10 homo-oligomers, can accommodate the translocation and secretion of such a diverse range of cargoes across different physiological and pathological contexts.

TMED10 is a member of the TMED protein family found in eukaryotic organisms. TMEDs are characterized as type I transmembrane proteins possessing a single transmembrane domain, and they display sequence similarities and share a comparable structural configuration (Aber et al., 2019; Pastor-Cantizano et al., 2016; Strating and Martens, 2009). In human, nine members of TMEDs have been reported (TMED1-7 & 9, 10, excluding TMED8 due to misannotation) (Strating et al., 2009). It is noteworthy that specific subsets of TMED proteins have been implicated in various cellular processes. These processes include but are not limited to the regulation of GPI-anchored proteins transport (Bonnon et al., 2010; Fujita et al., 2011; Muniz et al., 2000; Pastor-Cantizano et al., 2016; Schimmoller et al., 1995; Takida et al., 2008; Theiler et al., 2014),

ER-exit site/ERGIC/Golgi organization (Blum et al., 1999; Koegler et al., 2010; Mitrovic et al., 2008; Rojo et al., 2000; Yang et al., 2024), autophagy (Li et al., 2022; Shin et al., 2019), cell signaling (Connolly et al., 2013; Li et al., 2023; Liaunardy-Jopeace et al., 2014), membrane contact regulation (Anwar et al., 2022; Li et al., 2022), modulation of γ-secretase (Chen et al., 2006; Vetrivel et al., 2007), and Golgi-bypass trafficking (Park et al., 2022). Nevertheless, when it comes to the specific role of TMED10 in the regulation of cargo translocation, an intriguing question arises. It remains uncertain whether cargo translocation is a function unique to TMED10 or if it represents a common function shared among various TMED proteins.

## RESULTS

### TMEDs differentially regulate the secretion of active UcPS cargoes

We initiated an inquiry into the capacity of individual TMED proteins to facilitate the release of UcPS cargoes. In cell lines lacking TMED10 (TMED10-KO) and devoid of THU (Zhang et al., 2020), the levels of other TMEDs experienced significant reductions (Fig. S1, A and B). This outcome aligns with prior researches, which have consistently demonstrated diminished levels of TMEDs following TMED10 depletion due to mutual reliance on TMED10 for the stability of other TMEDs (Denzel et al., 2000; Fujita et al., 2011; Theiler et al., 2014). Given the dinimished basal levels of TMEDs, the TMED10-KO cells serve as a valuable model for examining the net impact of individual TMEDs on UcPS with single TMED expression.

Notably, transient expression (24 h) of individual V5-tagged TMED proteins resulted in distinct, cargo-specific effects on the release of various UcPS cargoes (the mature forms of interleukin(IL)-1ý, IL-1α, IL-33, IL-36α, IL-36RA, IL-37, Galectin-1, Galectin-3, Anexin-1A, HSPB5, and Tau) (Fig. 1, A and B; and Fig. S1, C-L). This effect appears specific to each TMED, as levels of other endogenous TMED proteins remained minimal throughout the experiment (Fig. S1 M). These findings suggest that individual TMEDs may play regulatory roles in UcPS with potential preferences for specific cargoes. As a control, the expression of ERGIC-localized single-membrane proteins, ERGIC53 or LMAN2, had no impact on the UcPS of IL-1β (Fig. S1 N).

**Figure 1.**
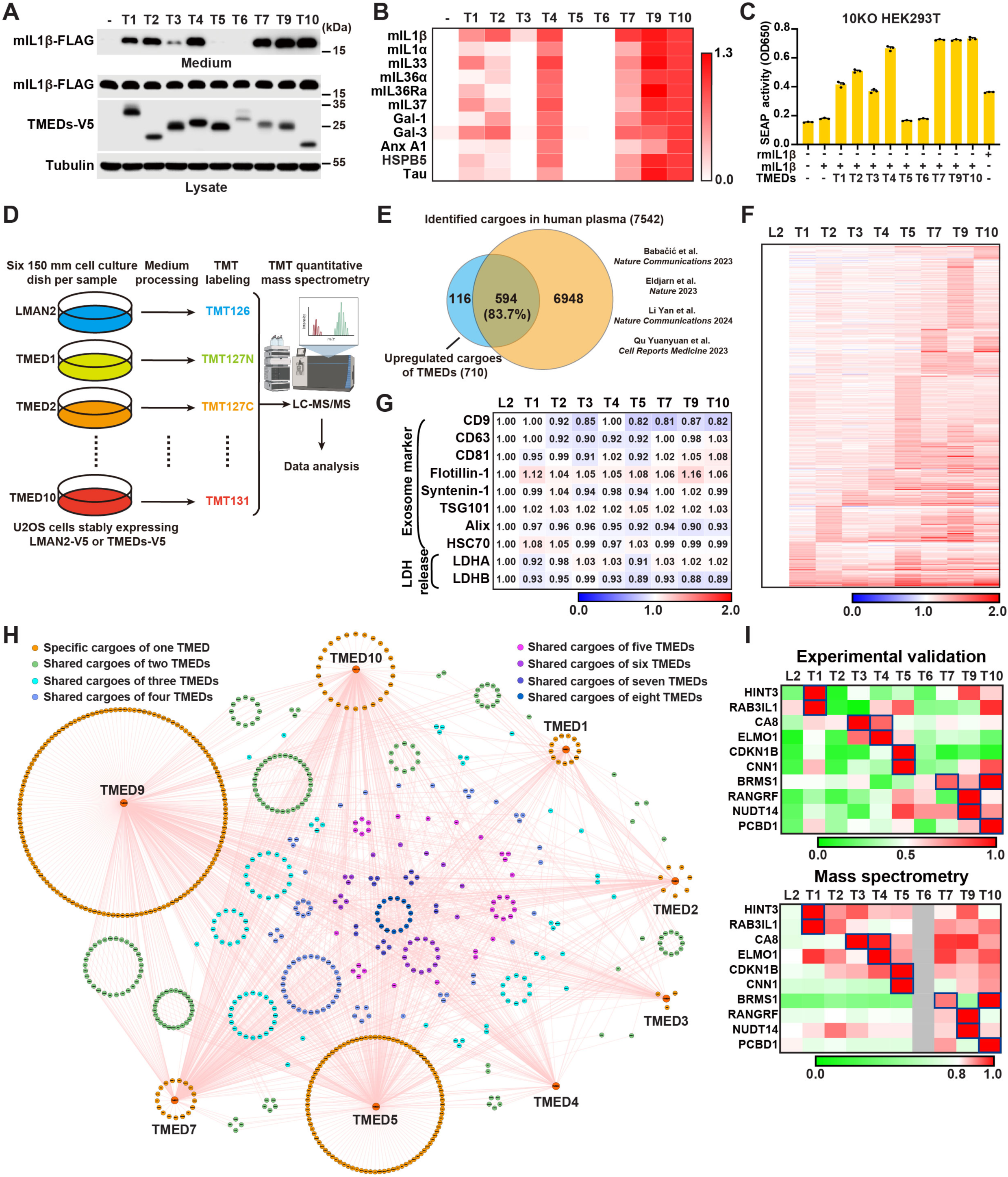
TMEDs differentially regulate the secretion of active UcPS cargoes. **(A)** Secretion of mIL-1β in TMED10-KO HEK293T cells with control or TMEDs (TMED1-7 & 9, 10) expression. The data are representative of three independent experiments. **(B)** Heatmap showing the secretion of indicated cargoes in TMED10-KO HEK293T cells with control or TMEDs (TMED1-7 & 9, 10) expression as shown in (A) and (Fig. S1, C-L). Relative level of cargo secretion was normalized to cargo level in cell lysates, and the TMED10 group was set as 1. **(C)** HEK-Blue IL-1β cells (InvivoGen) were treated with culture medium derived from TMED10-KO HEK293T cells with indicated protein expression, or with 0.1 μg/ml recombinant mIL-1β (rmIL-1β) as a positive control. Levels of SEAP indicating IL-1β activity in the medium were monitored using QUANTI-Blue (n = 3). **(D)** The quantitative secretomics workflow of U2OS cells with LMAN2-V5 or TMEDs-V5 (TMED1-5 & 7, 9, 10) stable expression. **(E)** Venn diagram depicting the overlap of upregulated cargoes identified in the secretome of TMEDs (TMED1-5 & 7, 9, 10) with proteins detected in human plasma. **(F)** Heatmap showing the secretome in U2OS cells with stable expression of LMAN2 or TMEDs (TMED1-5 & 7, 9, 10). Cargo secretion level in the LMAN2 group (control) was set as 1, and other sample groups were normalized relative to this control. The data are average of two independent experiments. **(G)** Relative level of indicated exosome markers and LDH release in the secretome of U2OS cells with LMAN2 or TMEDs (TMED1-5 & 7, 9, 10) stable expression. **(H)** Network diagram illustrating the relationships between different sets of upregulated UcPS cargoes of the TMEDs (TMED1-5 & 7, 9, 10). High-confidence upregulated UcPS cargoes unique to each TMED are arranged around the corresponding TMED, and shared cargoes are labeled with different colors denoting multiple combinations. **(I)** Heatmap showing the secretion of specific cargoes in U2OS cells with stable expression of LMAN2 or TMEDs (TMED1-7 & 9, 10), as measured by secretion assay (upper panel) shown in (Fig. S2, B-K) or mass spectrometry (lower panel). Relative level of cargo secretion was normalized to cargo expression in cell lysate and the group exhibiting the highest cargo secretion was set as 1. Blue boxes highlight the peak of specific TMED-enhanced secretion in both cellular secretion experiments and mass spectrometry.

Interestingly, when TMED10 expression was extended to 96 h in TMED10-KO cells, other TMED protein levels were restored, and typical TMED family functions—such as the facilitation of GPI-anchored protein transport and reestablishment of Golgi morphology—were also recovered (Fig. S1, O and P). This demonstrates that V5-tagged TMED10 retains its previously reported functional roles (Pastor-Cantizano et al., 2016). The data demonstrate that the V5-tag does not affect the function of TMED10 and therefore, we employed the V5-tagged TMED10 and similar tagged TMEDs for subsequent studies.

One possibility is that TMED expression may dump overexpressed and misfolded proteins similar to a previously reported unconventional secretory pathway termed misfolding-associated protein secretion (Lee et al., 2016). To ascertain the functionality of cargoes released by TMEDs, we conducted an IL-1β reporter assay (Zhang et al., 2020). The observed SEAP (secreted alkaline phosphatase) activity induced by IL-1β was in accordance with the levels of mIL-1β (mature interleukin-1β), present in the medium derived from cells expressing individual TMEDs (Fig. 1 C). This outcome suggests that TMEDs are involved in the regulation of the secretion of functional mIL-1β, and by extension, likely play a role in the secretion of other active forms of cargoes.

We replicated the IL-1β secretion and SEAP activity assays in wild-type 293T cells, yielding consistent trends in IL-1β secretion and corresponding SEAP activity (Fig. S1, Q and R). This suggests that cargo selectivity and the release of active cargoes by TMEDs are not exclusive to TMED10-KO cells. Utilizing TMED10-KO cells allows for the individual expression of each TMED (in short period (24 h)) without potential interference from other TMEDs reported previously, which could form hetero-oligomers and complicate the analysis (Pastor-Cantizano et al., 2016). Consequently, we primarily utilized TMED10 KO cells for subsequent experiments.

It is worth mentioning that TMED5 and TMED6 exhibited a limited capacity to promote UcPS with the cargoes tested (Fig. 1 B). The ineffectiveness of TMED5 is likely attributable to mis-localization, an aspect we delve into further in subsequent investigations (Fig. 5). In the case of TMED6, the lack of effect can be attributed to both mis-localization and inability to facilitate translocation in our current study (Fig. 2 and 5).

**Figure 2.**
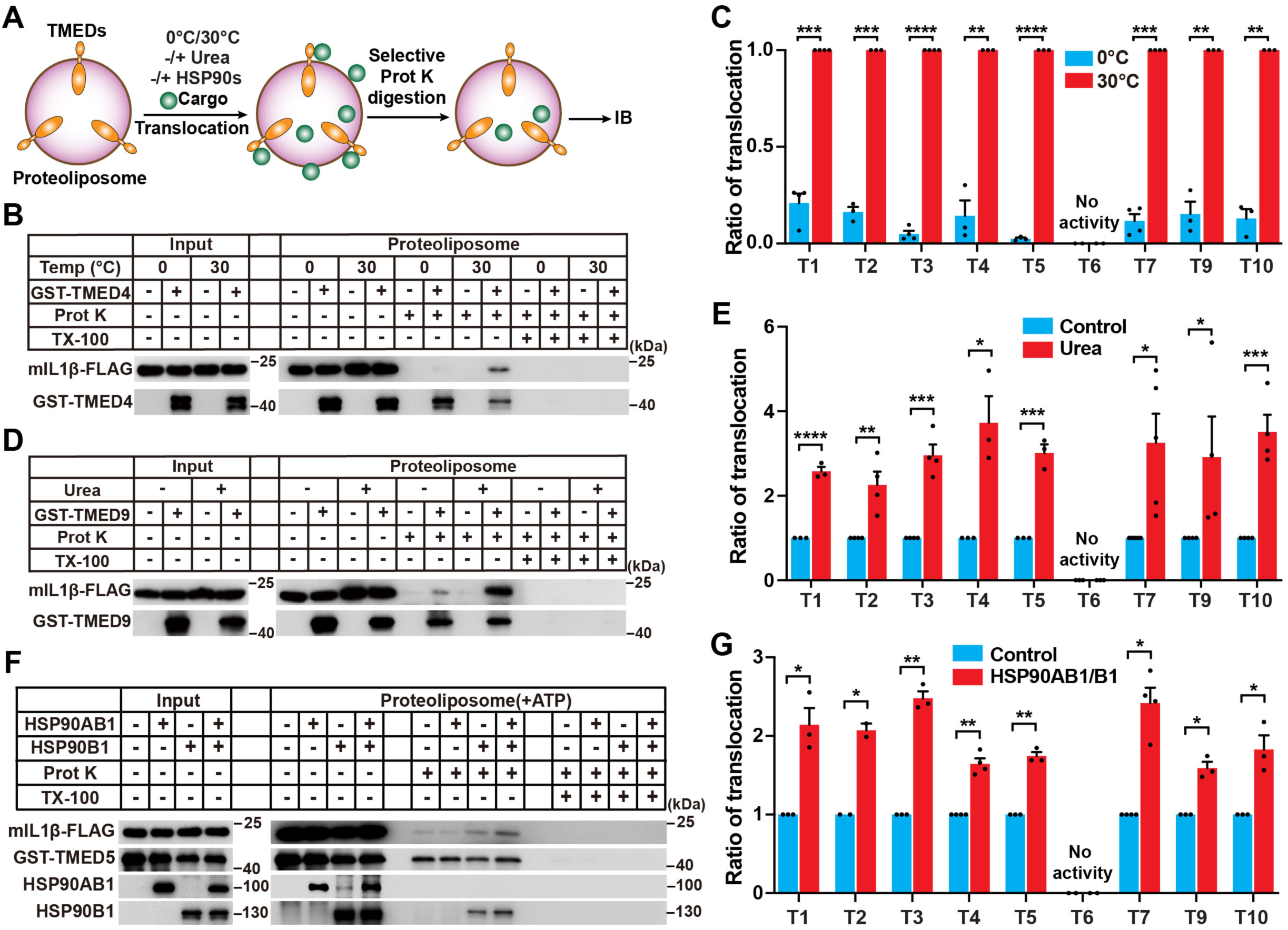
TMEDs directly translocate cargoes into liposomes. **(A)** Schematic diagram of the in vitro proteoliposome translocation assay. Briefly, proteoliposome containing GST-TMEDs (TMED1, 3-5, 7, 9, 10) or TMEDs (TMED2 & 6) were incubated with cargoes under various conditions (different temperatures, without or with 4 M urea pretreatment of cargo, and without or with HSP90s (HSP90AB1 outside and HSP90B1 inside the liposome)). Proteinase K digestion was used to assess the amount of mIL-1β translocated into the liposome. **(B)** In vitro translocation assay with control or GST-TMED4 proteoliposomes performed at 0°C and 30°C. **(C)** Quantification of the translocation efficiency of mIL-1β-FLAG (mean ± SEM) as shown in (B) and (Fig. S4 A); The relative level of translocation efficiency was normalized to TMEDs level and the group at 30°C was set as 1. p values were calculated by two-tailed t test (n ≥ 3). **(D)** In vitro translocation assay with control or GST-TMED9 proteoliposomes performed with mIL-1β without or with 4 M urea pre-treatment. **(E)** Quantification of the translocation efficiency of mIL-1β-FLAG (mean ± SEM) as shown in (D) and (Fig. S4 B); The relative level of translocation efficiency was normalized to TMEDs level and the group without 4 M urea pre-treatment was set as 1. p values were calculated by two-tailed t test (n ≥ 3). **(F)** GST-TMED5 proteoliposomes were generated in the presence or absence of HSP90B1. In vitro mIL-1β membrane translocation assay was performed with the proteoliposomes, ATP, and in the presence or absence of HSP90AB1. **(G)** Quantification of the translocation efficiency of mIL-1β-FLAG (mean ± SEM) in the presence or absence of HSP90s with ATP, as shown in (F) and (Fig. S4 C); The relative level of translocation efficiency was normalized to TMEDs level and the group without HSP90s was set as 1. p values were calculated by two-tailed t test (n ≥ 2). ns, non-significant; *P < 0.05; **P < 0.01; ***P < 0.001; ****P < 0.0001.

### TMEDs select and share cargoes in UcPS

In order to comprehensively assess the specificity of cargo selection, we conducted a secretome analysis using cells that express individual TMEDs (Fig. 1 D and Fig. S2 A), with the exception of TMED6 due to its loss of translocation activity. Using mass spectrometry, we identified 710 leaderless proteins whose secretion was upregulated by TMEDs (Table S1). Notably, 594 of these leaderless proteins have been previously detected in human plasma through proteomic studies (Babacic et al., 2023; Eldjarn et al., 2023; Li et al., 2024; Qu et al., 2023), suggesting that the majority of the identified secretory proteins are present extracellularly in humans (Fig. 1 E). Significantly, the expression of each TMED led to the enhanced secretion of distinct sets of UcPS cargoes, underscoring the specific cargo preferences of these proteins (Fig. 1 F). As controls, extracellular vesicle makers or LDH were not regulated by TMEDs indicating that the regulated release of cargoes are not exosomes or cell-death related (Fig. 1 G).

By utilizing network analysis in conjunction with a heatmap, we unveiled the distinct cargo profiles associated with each TMED. This analysis highlighted the presence of unique cargoes specific to individual TMEDs, as well as shared cargoes among multiple TMEDs, albeit often displaying varying degrees of enhanced secretion (Fig. 1, F and H). These observations suggest that each TMED is responsible for the management of its unique cargoes while also partially share cargoes with other TMEDs in the context of UcPS.

To validate this observed cargo selectivity, we selected ten UcPS cargoes that displayed clear specificity for one or two TMEDs in the secretome analysis and performed cellular secretion assays. The results from these cellular secretion assays revealed a consistent trend of TMED-enhanced secretion that aligns with the observations made through mass spectrometry (Fig. 1 I and Fig. S2, B-K). While manually validating all 710 cargoes may not be feasible, the findings from the cellular secretion assays at least provide evidence in support of the concept of cargo selectivity by each TMED in UcPS.

### TMEDs directly translocate cargoes into vesicles

One possible mechanism by which TMEDs facilitate UcPS is by destabilizing the plasma membrane, akin to the process observed in non-vesicular UcPS, as demonstrated with FGF2 release (Sparn et al., 2022). However, punicalagin, known to stabilize the plasma membrane (de Vasconcelos et al., 2019; Martin-Sanchez et al., 2016; Monteleone et al., 2018; Saeki et al., 2020) and effectively inhibit FGF2 release, did not impede the TMED-promoted release of UcPS cargoes (Fig. S3, A and B). This suggests that TMED-mediated UcPS may not rely on plasma membrane integrity loss, consistent with our prior findings demonstrating the independence of Gasdermin D-mediated pore formation (Zhang et al., 2020). Moreover, expression of multiple TMEDs in TMED10-KO cells resulted in elevated levels of membrane-localized UcPS cargoes, suggesting that TMED-mediated UcPS is vesicle-related and that TMEDs may facilitate cargo entry into secretory vesicles (Fig. S3, C-F).

Next, we aimed to investigate whether TMEDs possess the capability to translocate UcPS cargoes, a property previously demonstrated for TMED10 (Zhang et al., 2020). Multiple TMEDs have been shown to be glycosylated (Dominguez et al., 1998; Fullekrug et al., 1999). Consistently, TMED1, 4, 6, 7, and 9 showed a reduction of molecular weight after PNGase F treatment, indicating that these proteins are glycosylated (Fig. S3 G). However, mutation of the asparagine (the predicted glycosylation site) to glutamine did not affect the effects of these TMEDs on UcPS, indicating that glycosylation is not required for the function of TMEDs in UcPS (Fig. S3 H).

Since glycosylation does not impact the function of TMEDs on UcPS, we employed an E. coli expression system to purify GST-tagged TMEDs (Fig. S3 I). Subsequently, we reconstituted each TMED into proteoliposomes to assess their cargo translocation activity. Intriguingly, the purified TMEDs (specifically, TMED1-5, 7, 9, and 10) directly facilitated the translocation of mIL-1β, a cargo shared by multiple TMEDs (Fig. 1 A), into liposomes in a manner dependent on temperature (Fig. 2, A-C; and Fig. S4 A). Notably, pre-treatment of cargo with 4M urea augmented the translocation activity of TMEDs, suggesting that a partial cargo unfolding promotes the translocation process (Fig. 2, D and E; and Fig. S4 B). Furthermore, the translocation process was enhanced by the presence of two HSP90 proteins, HSP90B1 within the lumen and HSP90AB1 on the external side, in the presence of ATP (Fig. 2, F and G; and Fig. S4 C). Consequently, it can be inferred that TMEDs are proficient in translocating UcPS cargoes into liposomes, likely employing a mechanism involving cargo partial unfolding and the involvement of accessory factors such as HSP90s, the characters of which have been described in TMED10-mediated translocation (Zhang et al., 2020) and the molecular process is comparable to a previously identified translocation pathway of chaperone-mediated autophagy (Kaushik and Cuervo, 2018).

It’s worth noting that the majority of GST-tagged TMEDs displayed translocation activity, with the exception of TMED2, which required partial removal of the GST tag to restore its activity (Fig. S4). Notably, we observed no translocation activity from TMED6 under any of the tested conditions (Fig. 2, C, E and G). This observation aligns with the above observed absence of functionality for TMED6 in UcPS (Fig. 1 B).

### TMEDs translocate UcPS cargoes with selectivity

Our cellular secretion and secretome analysis indicate cargo selectivity of TMEDs. To determine if the selectivity originates from translocation, we aimed to establish a cell-free translocation system. This was necessitated by the potential disparities in translocation efficiency within the proteoliposome system, primarily arising from varying effects of the GST-tag on individual TMED functions. In this cell-free system, we transiently expressed (24 h) each TMED in TMED10-KO cells and combined the respective cellular membranes with cytosolic components, UcPS cargoes, and nucleotides to recreate the translocation process (Fig. 3 A).

**Figure 3.**
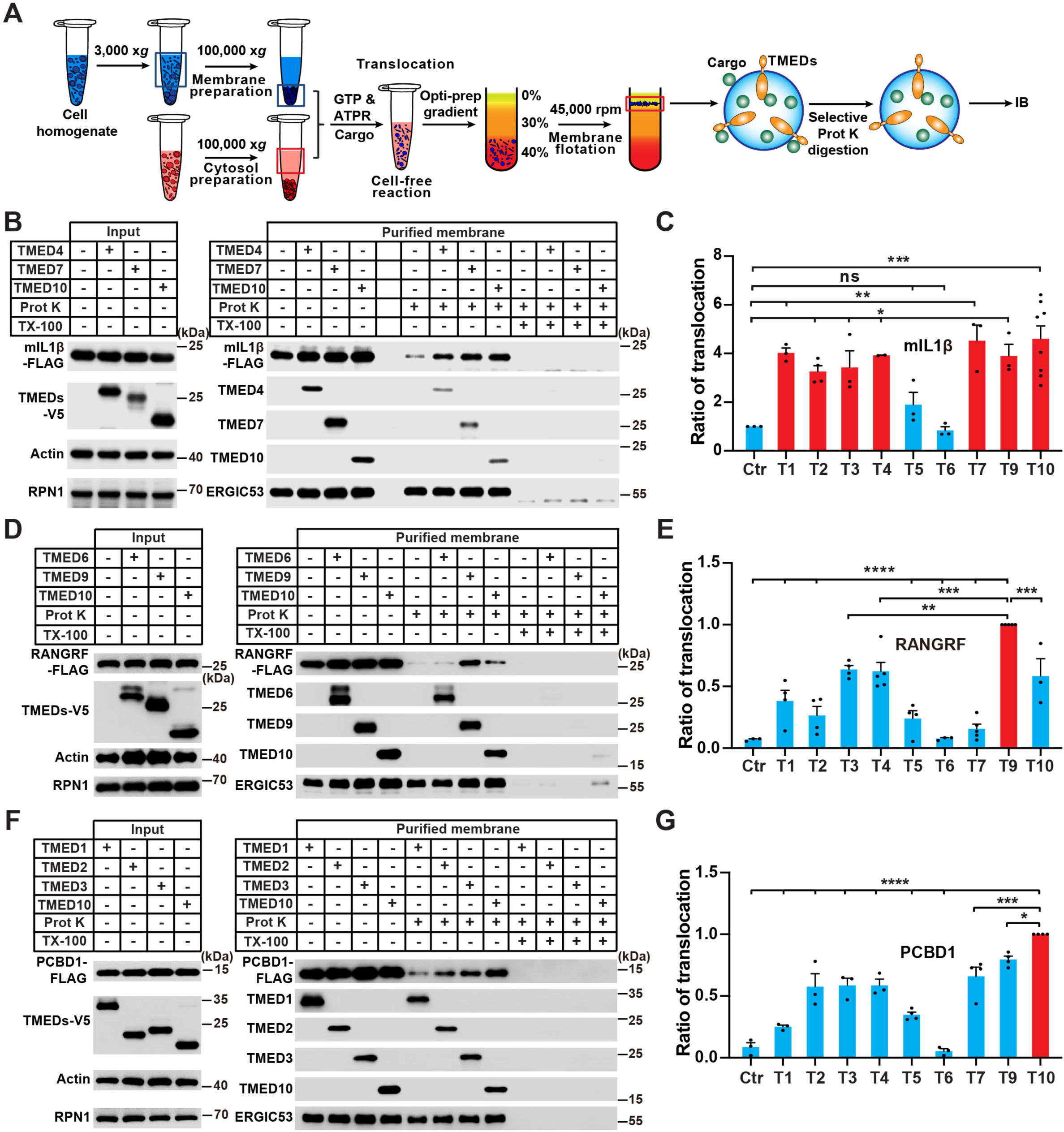
TMEDs directly translocate cargoes into vesicles with selectivity. **(A)** Schematic diagram of the cell-free translocation assay. Briefly, membranes from TMED10-KO HEK293T cells with control or TMEDs-V5 (TMED1-7 & 9, 10) expression were incubated with cargo, cytosol, ATPR & GTP to generate the cell-free translocation reaction. Membrane flotation and proteinase K digestion were then performed to determine the amount of membrane-incorporated cargo. ATPR: ATP regeneration system. **(B)** Cell-free translocation of mIL-1β using membranes from TMED10-KO HEK293T cells without or with TMEDs (TMED4, 7, 10) expression. **(C)** Quantification of the translocation efficiency of mIL-1β (mean ± SEM) without or with TMEDs (TMED1-7 & 9, 10) expression as shown in (B) and (Fig. S5, B and C), the relative level of mIL-1β translocation was normalized to the control group; p values were calculated by one-way analysis of variance (ANOVA) (n ≥ 2). **(D)** Cell-free translocation of RANGRF using membranes from TMED10-KO HEK293T cells without or with TMEDs (TMED6, 9, 10) expression. **(E)** Quantification of the translocation efficiency of RANGRF (mean ± SEM) without or with TMEDs (TMED1-7 & 9, 10) expression as shown in (D) and (Fig. S5, D and E), the relative level of RANGRF translocation was normalized to the TMED9 group; p values were calculated by one-way ANOVA (n ≥ 3). **(F)** Cell-free translocation of PCBD1 using membranes from TMED10-KO HEK293T cells with TMEDs (TMED1-3 & 10) expression. **(G)** Quantification of the translocation efficiency of PCBD1 (mean ± SEM) without or with TMEDs (TMED1-7 & 9, 10) expression as shown in (F) and (Fig. S5, F and G), the relative level of PCBD1 translocation was normalized to the TMED10 group; p values were calculated by one-way ANOVA (n ≥ 3). ns, non-significant; *P < 0.05; **P < 0.01; ***P < 0.001; ****P < 0.0001.

TMED10, when expressed and localized on the intracellular membrane fraction, promoted the entry of mIL-1β into the membrane compartment, and the reaction was further activated by nucleotides (Fig. S5 A). Additionally, several TMEDs (TMED1-4, 7, 9, and 10) promoted the translocation of mIL-1β into the membrane compartment (Fig. 3, B and C; and Fig. S5, B and C), consistent with the outcomes observed in the proteoliposome-based translocation system. TMED5 exhibited largely reduced activity likely due to its location out of the ERGIC when transiently expressed (additional data shown in Fig. 5). TMED6 was consistently inactive in the cell-free assay.

We next employed the cell-free translocation system to determine the selectivity of TMEDs translocation using TMED9 and TMED10 as examples. As shown above, the secretion of cargoes RANGRF and PCBD1 were most efficiently promoted by TMED9 and TMED10, respectively (Fig. 1 I). Consistent with the selectivity of secretion, the translocation of RANGRF was mostly enhanced by TMED9 and the entry of PCBD1 into the membrane was predominantly promoted by TMED10 (Fig. 3, D-G; and Fig. S5, D-G). Therefore, the data suggest that TMEDs-mediated specificity of cargo translocation may be a major determinant of cargo selectivity in TMEDs-facilitated UcPS.

### The cytoplasmic tail of TMEDs is a determinant of cargo selection

Our previous work demonstrated the critical involvement of the cytoplasmic tail (CT) of TMED10 in cargo binding and translocation (Zhang et al., 2020). We explored whether the CT contributes to the differential cargo selection between TMED9 and TMED10. Using a Duo-link proximity ligation assay, TMED9 and TMED10 exhibited preferential binding to RANGRF and PCBD1, respectively (Fig. 4, A-D). Deletion of the CT abolished the interaction of TMED9 or TMED10 with their cargoes, while swapping CTs resulted in switched cargo binding selectivity (Fig. 4, A-D). This role was confirmed in secretion and cell-free translocation assays, where CT determined cargo selectivity (Fig. 4, E-H). Nonetheless, the potential involvement of other domains, such as the GOLD or coiled-coil domains, cannot be overlooked. These domains regulate TMED oligomerization, potentially impacting cargo translocation dynamics and efficiency.

**Figure 4.**
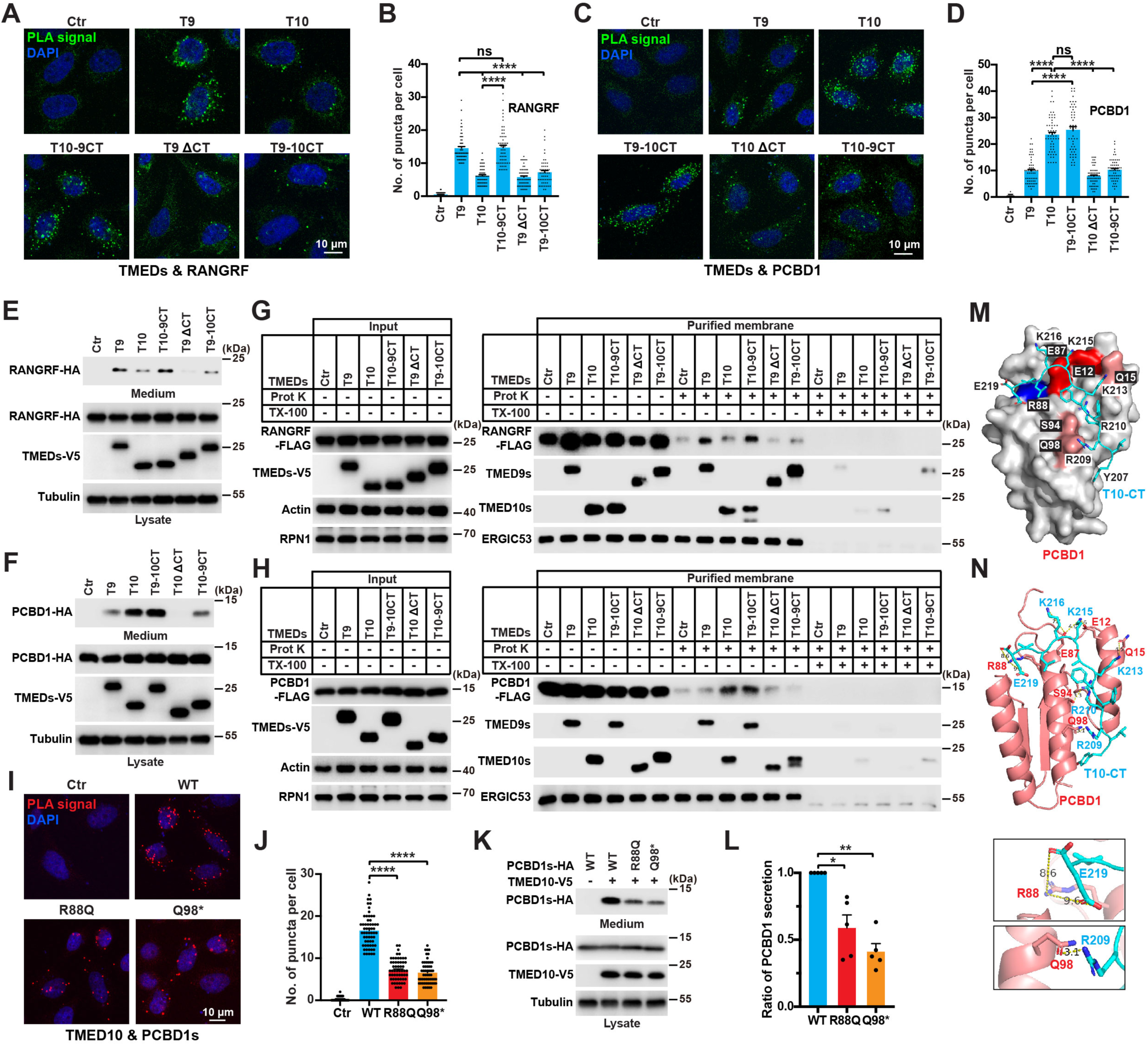
The cytoplasmic tail of TMEDs contributes to cargo selection. **(A and B)** Duolink PLA assay performed with HeLa cells expressing empty vector (Ctr), TMED9-V5 (T9), TMED10-V5 (T10), TMED10-TMED9CT (cytoplasmic tail, T10-9CT), TMED9ΔCT (T9ΔCT), TMED9-TMED10CT (T9-10CT) respectively and RANGRF-FLAG to test their interactions (A). The puncta of duolink signal area per cell (mean ± SEM) was quantified in (B); p values were calculated by one-way ANOVA (> 50 cells from three independent experiments). Scale bar, 10 μm. **(C and D)** Duolink PLA assay performed with HeLa cells expressing empty vector (Ctr), TMED9-V5 (T9), TMED10-V5 (T10), TMED9-TMED10CT(T9-10CT), TMED10ΔCT (T10ΔCT), TMED10-TMED9CT (T10-9CT) respectively and PCBD1-FLAG to test their interactions (C). The puncta of duolink signal area per cell (mean ± SEM) was quantified in (D); p values were calculated by one-way ANOVA (> 50 cells from three independent experiments). Scale bar, 10 μm. **(E and F)** Secretion of RANGRF (E) or PCBD1 (F) in TMED10-KO HEK293T cells with control or indicated TMED variants expression. The data are representative of three independent experiments. **(G and H)** Cell-free translocation of RANGRF (G) or PCBD1 (H) using membranes from TMED10-KO HEK293T cells without or with indicated TMED variants expression. The data are representative of three independent experiments. (I) Duolink PLA assay performed with HeLa cells expressing empty vector (Ctr), PCBD1 wild type (WT), or disease-associated mutations R88Q and Q98* respectively and TMED10-V5 to test their interaction. Scale bar, 10 μm. **(J)** Quantification of the puncta of duolink signal area per cell (mean ± SEM) as shown in (I); p values were calculated by one-way ANOVA (> 50 cells from three independent experiments). **(K)** Secretion of PCBD1 WT, disease-associated mutations R88Q and Q98* in TMED10-KO HEK293T cells with control or TMED10-V5 expression. **(L)** Quantification of the PCBD1 WT, disease-associated mutations R88Q and Q98* secretion (mean ± SEM) as shown in (K); p values were calculated by one-way ANOVA (n = 5). **(M)** Surface overview showing multiple charge interactions and hydrogen bonds between PCBD1 and T10-CT (TMED10 cytoplasmic tail) which was predicted by AlphaFold 3. **(N)** Upper, Ribbon diagram of the interaction between PCBD1 and T10-CT. PCBD1 is colored in pink and T10-CT is colored in cyan. Lower, two close-up views of the ribbon diagram showing the charge interactions of PCBD1-R88 with TMED10-E219, and hydrogen bond between PCBD1-Q98 and TMED10-R209. ns, non-significant; *P < 0.05; **P < 0.01; ***P < 0.001; ****P < 0.0001.

**Figure 5.**
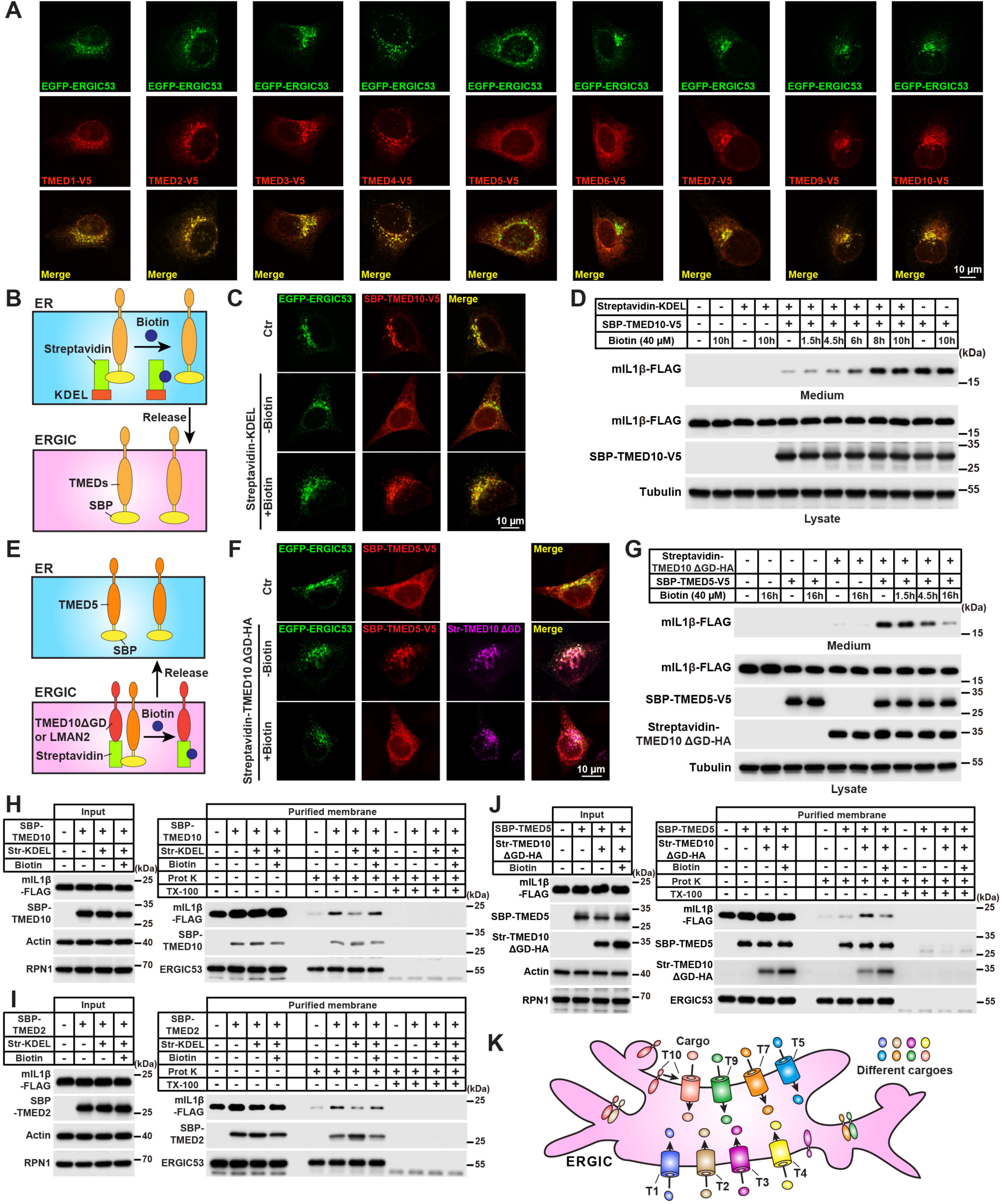
ERGIC localization determines TMED activity. **(A)** Immunofluorescence of U2OS cells co-expressing EGFP-ERGIC-53 and TMEDs-V5 with anti-V5 antibodies. Scale bar, 10 μm. **(B)** Schematic diagram of the RUSH system to retain ERGIC-localized TMEDs (TMED1-4 & 7, 9, 10) to the ER. TMEDs fused with a streptavidin binding peptide (SBP) are retained in the ER via binding to streptavidin (Str) connected to a KDEL sequence. Upon biotin addition, TMEDs were released and trafficked to ERGIC. **(C)** Immunofluorescence of TMED10-KO HeLa cells co-expressing EGFP-ERGIC-53 and SBP-TMED10-V5, without or with Str-KDEL expression, and treated without or with 40 μM biotin for 8 h. Scale bar, 10 μm. **(D)** Secretion assay of mIL-1β in TMED10-KO HEK293T cells combined with RUSH system. Cells were co-transfected without or with mIL-1β-FLAG, Str-KDEL, and SBP-TMED10-V5 and treated without or with biotin at the indicated concentration and time points. The data are representative of three independent experiments. **(E)** Schematic diagram of the RUSH system to retain ER-localized TMED5 to the ERGIC. TMED5 fused with SBP was retained via binding to streptavidin connected to TMED10ΔGOLD or LMAN2, two ERGIC-localized membrane protein deficient in UcPS. Upon biotin addition, TMED5 was released and trafficked back to the ER. **(F)** Immunofluorescence of TMED10-KO HeLa cells co-expressing EGFP-ERGIC-53 and SBP-TMED5-V5, without or with Str-TMED10ΔGOLD-HA expression, and treated without or with 40 μM biotin for 16 h. Scale bar, 10 μm. **(G)** Secretion assay of mIL-1β in TMED10-KO HEK293T cells combined with RUSH system. Cells were co-transfected without or with mIL-1β-FLAG, Str-TMED10ΔGOLD-HA, SBP-TMED5-V5 and treated without or with biotin at the indicated concentration and time points. The data are representative of three independent experiments. **(H and I)** Cell-free translocation assay of mIL-1β in TMED10-KO HEK293T cells combined with RUSH system. Cells were transfected without or with Str-KDEL and SBP-TMED10-V5 (H) or SBP-TMED2-V5 (I) and treated without or with 40 μM biotin for 16 h. The data are representative of three independent experiments. **(J)** Cell-free translocation assay of mIL-1β in TMED10-KO HEK293T cells combined with RUSH system. Cells were transfected without or with Str-TMED10ΔGOLD-HA and SBP-TMED5-V5 and treated without or with 40 μM biotin for 16 h. The data are representative of three independent experiments. **(K)** A model showing diversified UcPS cargo translocation mediated by TMEDs.

To investigate a potential pathological link, we analyzed PCBD1, mutations of which are associated with metabolic disorders such as early onset diabetes, hypomagnesemia, and hyperphenylalaninemia (Ferre et al., 2014; Simaite et al., 2014; Thony et al., 1998a; Thony et al., 1998b). We introduced disease-associated mutations into PCBD1 and assessed their effects, the R88Q and Q98* (with residues 98-104 deleted) which remained stable when expressed in HEK293T. Further analysis revealed that these mutants had reduced interactions with TMED10 and impaired unconventional protein secretion (UcPS), suggesting a connection between PCBD1 UcPS and the aforementioned diseases (Fig. 4, I-L). Structural modeling of the PCBD1-TMED10-CT interaction shows that residues R88 and Q98 are crucial for several charged interactions and a hydrogen bond with corresponding residues (E219 and R209) of TMED10-CT, explaining the reduced binding affinity and subsequent impairment in UcPS observed with these mutants (Fig. 4, M and N).

Collectively, these data suggest that the CT region of TMED proteins serves as a critical determinant for cargo selectivity in TMED-mediated translocation and unconventional secretion. Impairments in CT-mediated cargo selection may contribute to the pathogenesis of specific metabolic disorders.

### ERGIC localization determines UcPS mediated by TMEDs

Among the TMED proteins, TMED1-4, 7, 9, and 10 consistently exhibit activity in both UcPS and the translocation process, while TMED6 consistently remains inactive in these assays. In the case of TMED5, we observed variable activity across different experiments. When transiently expressed in TMED10-KO cells, TMED5 had no discernible effect on UcPS and cargo translocation in the cell-free assay (Fig. 1 B and Fig. 3 C) but was active in the translocation process using proteoliposomes (Fig. 2, C, E and G).

To address this inconsistency, we conducted an analysis of the localization of TMED5 and other TMEDs. Upon transient expression, the majority of the TMEDs (specifically, TMED1-4, 7, 9, and 10) were found within the ERGIC, whereas TMED5 (and TMED6) were localized in the ER (Fig. 5 A). These observations suggest that the localization of TMED may play a pivotal role in determining its function in UcPS, with activation occurring when it is situated in the ERGIC but not when localized in the ER.

To assess whether ERGIC localization is essential for activating the UcPS function of TMED5 and other TMEDs, we leveraged the fact that TMEDs are known to shuttle between the ER and the ERGIC via membrane trafficking (Dominguez et al., 1998; Gommel et al., 1999; Pastor-Cantizano et al., 2016). We employed a Retention Using Selective Hooks (RUSH) system to retain the ERGIC-localized TMEDs (TMED1-4, 7, 9, 10) within the ER, achieved by employing an ER-hook (Streptavidin-KDEL) (Boncompain et al., 2012). The TMEDs tagged with SBP (streptavidin binding peptide) can be released from these hooks through biotin, which competes against the SBP-hook interaction. The retention of ERGIC-localized TMEDs within the ER nullified their capacity to enhance UcPS, and this functionality was subsequently restored upon biotin treatment (Fig. 5, B-D and Fig. S6, A-L).

Conversely, for the transiently expressed TMED5 located within the ER, re-localizing it to the ERGIC using an ERGIC-hook (Streptavidin-TMED10ΔGOLD) led to a UcPS-promoting function of TMED5. This function was then reversed when TMED5 was re-localized back to the ER (Fig. 5, E-G). We then utilized an alternative ERGIC-targeting strategy (Streptavidin-LMAN2), which similarly recruits TMED5 to the ERGIC, thereby activating its UcPS function (Fig. S6, M and N). These findings support the conclusion that the ERGIC localization of TMEDs, including TMED1-5, 7, 9, and 10, is pivotal in activating their capability to facilitate vesicle-mediated unconventional protein secretion.

It is noteworthy that TMED proteins may also partially localize to the Golgi. To ascertain whether translocation mediated by TMEDs occurs at the Golgi rather than the ERGIC, we employed an O-glycosylation assay, in which an O-GalNAcylation motif (GATGAGAGAGTTPGPG (de Las Rivas et al., 2019)) was inserted into mIL-1β. As a positive control, mIL-1β containing both a signal peptide and the O-GalNAcylation motif underwent ER-Golgi trafficking, exhibiting a mobility shift in both the medium and lysate fractions indicative of O-GalNAcylation, which was confirmed through O-glycoprotease digestion (Fig. S6, O and P). In contrast, mIL-1β with only the O-GalNAcylation motif was released via UcPS and did not undergo O-GalNAcylation, as evidenced by the absence of mobility shift and resistance to O-glycoprotease digestion (Fig. S6, O and P). Thus, our findings indicate that TMED-mediated translocation occurs at the ERGIC rather than the Golgi.

### ERGIC localization is required for TMED-mediated translocation

The data presented above highlight the necessity of ERGIC localization for TMED-mediated UcPS. We next sought to investigate the regulation of translocation and its relationship with ERGIC localization. Since the translocation assay conducted with proteoliposomes does not account for the subcellular localization of TMEDs, we employed the cell-free translocation system that utilized intracellular membranes to examine the connection between ERGIC localization and TMED-mediated translocation. As mentioned above, multiple TMEDs localized in the ERGIC (specifically, TMED1-4, 7, 9, and 10) promoted the translocation of mIL-1β into the membrane compartment, while the ER-localized TMED5 and TMED6 displayed significantly reduced activity (Fig. 3 C). These results align with the findings of the secretory assay detailed earlier (Fig. 1 B).

We integrated the cell-free translocation system with the RUSH method to assess the necessity of ERGIC localization for TMED-mediated translocation. Membrane fractions of TMED10 or TMED2, both of which typically reside within the ERGIC, exhibited compromised TMED-promoted translocation when retained within the ER. Conversely, releasing these membrane fractions to the ERGIC restored translocation activity (Fig. 5, H and I). In a reciprocal experiment, membrane fractions of TMED5, an ER-localized TMED, displayed an enhanced translocation function when anchored to the ERGIC. However, when TMED5 were retrieved back to the ER, the membrane exhibited reduced translocation activity (Fig. 5 J). Hence, these consistent findings underscore the crucial role of ERGIC localization in activating TMED-mediated translocation within the context of UcPS.

Collectively, the data indicate that ERGIC localization is a crucial prerequisite for the activation of TMED proteins as translocators involved in UcPS cargo translocation. This process subsequently initiates a Golgi-bypass mechanism, which is facilitated by ERGIC-localized Rab proteins (Sun et al., 2024). Therefore, TMED family can function as translocators, selectively transporting cargoes at the ERGIC and facilitating the release of diverse UcPS cargoes (Fig. 5 K).

### Switch of oligomerization regulates TMEDs function in translocation and UcPS

The TMED protein family exhibits four distinct subfamilies denoted as α (comprising TMED4 and TMED9), β (TMED2), γ (TMED1, TMED3, TMED5, TMED6, and TMED7), and δ (TMED10) (Aber et al., 2019; Pastor-Cantizano et al., 2016; Schuiki and Volchuk, 2012; Strating and Martens, 2009). These subfamily members are known to assemble into hetero-tetrameric complexes, containing one representative from each of the α, β, γ and δ subfamilies. These complexes primarily function in the intracellular transport of cargoes, such as GPI-anchored proteins, from the endoplasmic reticulum (ER) to the Golgi apparatus (Bonnon et al., 2010; Fujita et al., 2011; Muniz et al., 2000; Pastor-Cantizano et al., 2016; Schimmoller et al., 1995; Takida et al., 2008; Theiler et al., 2014; Yang et al., 2024). Furthermore, the presence of the hetero-tetramer is crucial for maintaining the stability of TMED proteins (Denzel et al., 2000; Fujita et al., 2011; Marzioch et al., 1999; Theiler et al., 2014). Intriguingly, label-free mass spectrometry quantification experiments showed that the four TMED subfamilies are not present in an exact 1:1:1:1 ratio in two distinct cell lines (Fig. 6, A-C). Furthermore, a comprehensive proteomic study of hundreds of human cancer cell lines across multiple cancer types confirms this imbalance (Fig. 6 D) (Goncalves et al., 2022). These data strongly suggest that TMEDs do not exclusively exist as hetero-tetramers, although the stability of individual subfamily members relies on interactions with one another. This observation is further supported by size exclusion analyses, which demonstrated the existence of multiple homo- and hetero-oligomeric forms (or even a monomer in complex with other proteins), in addition to the tetrameric structure, within the cellular context (Fig. 6 E), which is consistent with previous studies (Dvela-Levitt et al., 2019; Jenne et al., 2002; Maldutyte et al., 2025; Marzioch et al., 1999; Mota et al., 2022; Nagae et al., 2016; Nagae et al., 2017; Xiao et al., 2024).

**Figure 6.**
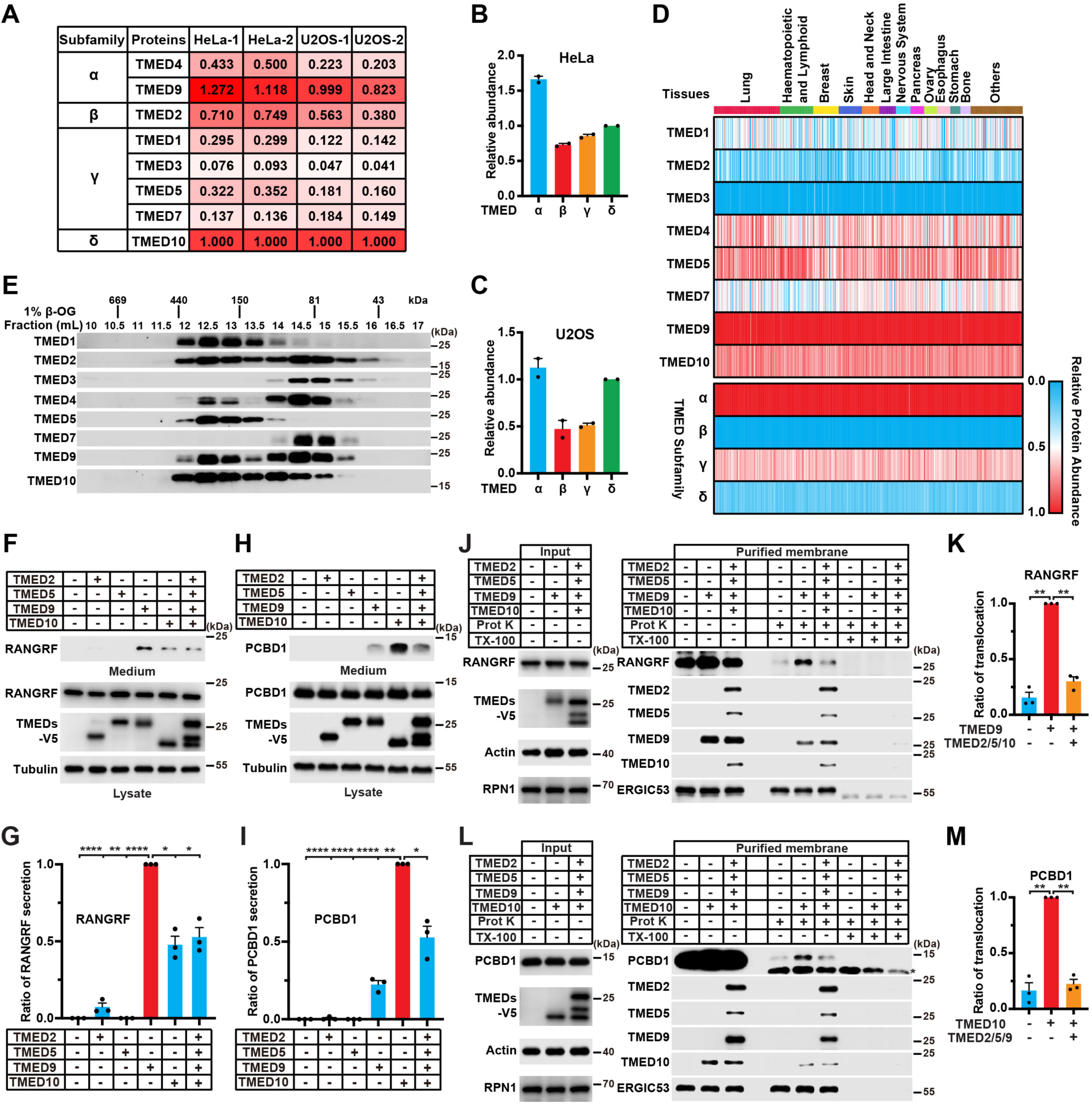
Switch of oligomerization regulates TMEDs function. **(A)** Label-free mass spectrometry quantification showing the relative amount of endogenous TMEDs (TMED1-5 & 7, 9, 10) in HeLa and U2OS cells with two replicates respectively. TMED6 was undetectable in this two cell lines. **(B and C)** Quantification of the relative abundance of four TMED subfamilies in HeLa (B) and U2OS (C) cells as shown in (A). **(D)** Heatmap showing the relative abundance of TMEDs (TMED1-5 & 7, 9, 10) and four subfamilies in human cell lines from multiple tissues. **(E)** Gel filtration assay analyzing the endogenous TMEDs (TMED1-5 & 7, 9, 10). HeLa cells were lysed and then applied to a Superdex 200 HR 10/30 column in 1% β-OG buffer. **(F and H)** Secretion of RANGRF (F) or PCBD1 (H) in TMED10-KO HEK293T cells with control or individual expression of four TMED subfamily members (TMED2, 5, 9, 10) or simultaneous expression to form hetero-tetramer. **(G and I)** Quantification of the RANGRF (G) or PCBD1 (I) secretion (mean ± SEM) as shown in (F) and (H); p values were calculated by one-way ANOVA (n = 3). **(J)** Cell-free translocation of RANGRF using membranes from TMED10-KO HEK293T cells expressing TMED9 alone or with three other subfamily members (TMED2, 5, 10) co-expression. **(K)** Quantification of the translocation efficiency of RANGRF (mean ± SEM) as shown in (J), the relative level of RANGRF translocation was normalized to the TMED9 group; p values were calculated by one-way ANOVA (n = 3). **(L)** Cell-free translocation of PCBD1 using membranes from TMED10-KO HEK293T cells expressing TMED10 alone or with three other subfamily members (TMED2, 5, 9) co-expression. **(M)** Quantification of the translocation efficiency of PCBD1 (mean ± SEM) as shown in (L), the relative level of PCBD1 translocation was normalized to the TMED10 group; p values were calculated by one-way ANOVA (n = 3). ns, non-significant; *P < 0.05; **P < 0.01; ***P < 0.001; ****P < 0.0001.

Our previous findings indicate that TMED10 self-assembles into homo-oligomers, forming a protein translocator that facilitates protein translocation (Zhang et al., 2020). To further investigate the regulatory mechanisms underlying UcPS, we examined the interplay between homo-oligomerization (identified in this study) and hetero-tetramerization (as previously reported) in modulating UcPS and cargo translocation. We selected two cargoes specific to TMED9 and TMED10, RANGRF and PCBD1, to assess their secretion and translocation. Remarkably, TMED9 or TMED10 alone promoted efficient secretion and translocation of RANGRF or PCBD1. In contrast, co-expression of TMED9 or TMED10 with three other subfamily members led to hetero-tetramer formation, which significantly inhibited cargo secretion and translocation (Fig. 6, F-M). These results suggest that hetero-tetramerization impairs the ability of TMED proteins to function as effective translocators for UcPS cargo.

### Localization-specific formation of two different TMED oligomers

TMEDs cycle between the ER and the ERGIC. While our previous work demonstrate that ERGIC localization is required for the function of TMED-mediated translocation and UcPS. We next employed a previous established membrane fractionation approach to isolate ER and ERGIC-enriched fractions to determine preferential form of TMED oligomer at the ER and the ERGIC. An ERGIC-enriched fraction (L) and an ER-enriched fraction (P) was prepared (Ge et al., 2013). Co-immunoprecipitation analysis focusing on TMED10 self-association or association with other subfamily members was performed to indicate homo- or hetero-oligomer formation. Intriguingly, we observed a higher propensity for TMED10 self-association (the homo-oligomer) within the ERGIC-enriched fraction as compared to the ER-enriched fraction (Fig. 7, A-C). Conversely, the association of TMED10 with other subfamily members (the hetero-oligomer) exhibited the opposite pattern (Fig. 7, A-C).

**Figure 7.**
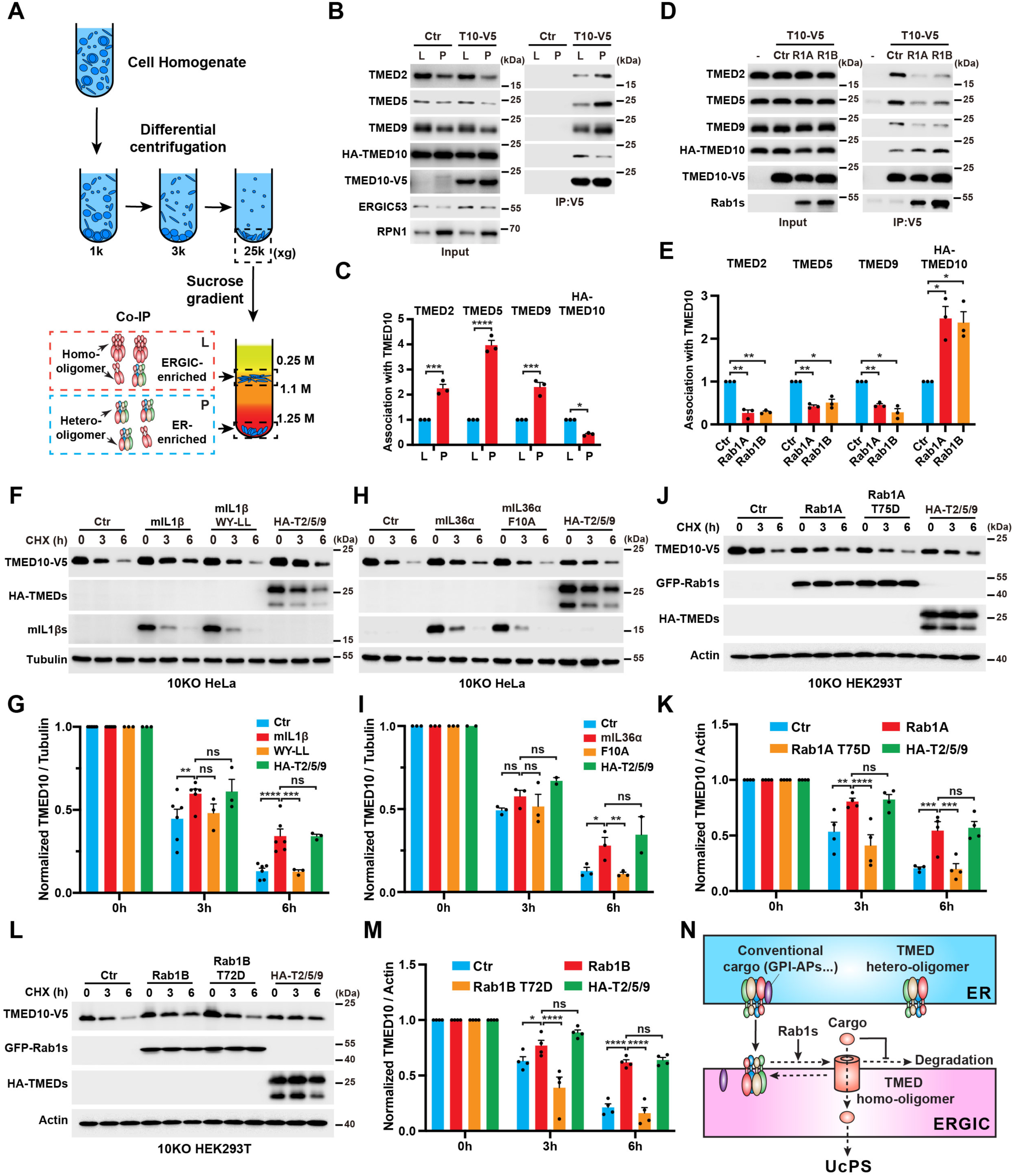
Localization regulates TMED oligomerization states and UcPS cargoes stabilize the TMED homo-oligomer. **(A)** Schematic diagram of membrane fractionation using differential centrifugation and sucrose gradient centrifugation to separate ERGIC-enriched L fraction and ER-enriched P fraction followed by co-immunoprecipitation. **(B)** Co-IP (Co-immunoprecipitation) assay performed with the ERGIC-enriched fraction (L) and the ER-enriched fraction (P) extracted from HEK293T cells expressing HA-TMED10, without or with TMED10-V5 co-expression using anti-V5 agarose. **(C)** Quantification of the association efficiency of TMED10-V5 with other subfamily members (TMED2, 5, 9) or HA-TMED10 (mean ± SEM) as shown in (B); p values were calculated by two-way ANOVA (n = 3). **(D)** Co-IP assay was performed with HEK293T cells co-expressing TMED10-V5 and HA-TMED10 without or with GFP-Rab1A (R1A) or GFP-Rab1B (R1B) using anti-V5 agarose. **(E)** Quantification of the association efficiency of TMED10-V5 with other subfamily members (TMED2, 5, 9) or HA-TMED10 (mean ± SEM) as shown in (D); p values were calculated by two-way ANOVA (n = 3). **(F)** Turnover of TMED10-V5 in CHX chase assay without or with mIL-1β-FLAG or its UcPS-deficient mutant mIL-1β-WY-LL or three other subfamily members (HA-TMED2, 5, 9) expression in TMED10 KO HeLa cells. **(G)** Quantification of normalized TMED10-V5 (mean ± SEM) as shown in (F), the relative level of TMED10-V5 was normalized to Tubulin and the 0 h control group was set as 1; p values were calculated by two-way ANOVA (n ≥ 3). **(H)** Turnover of TMED10-V5 in CHX chase assay without or with mIL-36α, mIL-36α-F10A and HA-TMED2, 5, 9 expression in TMED10 KO HeLa cells. **(I)** Quantification of normalized TMED10-V5 (mean ± SEM) as shown in (H); p values were calculated by two-way ANOVA (n ≥ 2). **(J)** Turnover of TMED10-V5 in CHX chase assay without or with RAB1A, RAB1A UcPS-deficient mutant T75D and HA-TMED2, 5, 9 expression in TMED10 KO HEK293T cells. **(K)** Quantification of normalized TMED10-V5 (mean ± SEM) as shown in (J); p values were calculated by two-way ANOVA (n = 4). **(L)** Turnover of TMED10-V5 in CHX chase assay without or with RAB1B, RAB1B UcPS-deficient mutant T72D and HA-TMED2, 5, 9 expression in TMED10 KO HEK293T cells. **(M)** Quantification of normalized TMED10-V5 (mean ± SEM) as shown in (L); p values were calculated by two-way ANOVA (n = 4). **(N)** Model showing two distinct populations of TMED oligomers exist in cellular trafficking pathway. TMED hetero-oligomers play a role in conventional cargo (e.g. GPI-anchored proteins) transport between ER and ERGIC, while TMED homo-oligomers mediate unconventional secretion cargo translocation at the ERGIC. UcPS cargo stabilizes the homo-oligomeric structures, which subsequently facilitates translocation. ns, non-significant; *P < 0.05; **P < 0.01; ***P < 0.001; ****P < 0.0001.

Previously, we identified ERGIC-localized Rab1A and Rab1B as enhancers of TMED10 homo-oligomerization (Sun et al., 2024). To further investigate their role in oligomerization balance, we examined the effects of Rab1A and Rab1B on TMED10 assembly. Interestingly, both Rab1A and Rab1B promoted TMED10 self-association while reducing its interaction with other TMEDs, indicating a shift toward homo-oligomerization rather than hetero-oligomerization (Fig. 7, D and E). These findings suggest that TMED oligomerization state is dynamically regulated by its subcellular localization (ERGIC vs. ER) and ERGIC-localized Rab1s, enabling distinct functional roles in different cellular compartments.

### UcPS cargoes stabilize the TMED homo-oligomer

Each TMED subfamily relies on the presence of other subfamily members for stability. Notably, in TMED10-KO cells, transient expression of TMED10 alone resulted in its degradation within approximately six hours (Fig. 7, F and G). However, this degradation was mitigated by the co-expression of other TMED members, which facilitated hetero-tetramer formation, aligning with previous findings that underscore the role of hetero-tetramers in maintaining TMED stability (Fig. 7, F and G).

Co-expression of UcPS cargoes, such as mIL-1β or mIL-36, significantly increased the stability of TMED10 without the requirement for hetero-tetramer formation (Fig. 7, F-I; and Fig. S7, A-G). Notably, the stabilizing effect exerted by the cargoes was comparable to the stability conferred by hetero-tetramer formation in HeLa cells, and moderately lower in HEK293T cells (Fig. 7, F-I; and Fig. S7, A-G). This suggests that the cargo-induced stabilization of TMED10 is on par with hetero-tetramer formation, indicating that cargo binding may be sufficient to sustain the long-term stability of a single TMED10 unit. This finding uncovers a previously unrecognized mechanism for the maintenance of single TMED10 stability, which may also extend to other TMED family members.

In addition, this stabilizing effect was highly specific, as it was abolished when the cargo contained a previously reported (Zhang et al., 2020) mutated signal sequence that impaired UcPS, thereby nullifying the stabilization (Fig. 7, F-I; and Fig. S7, A-G). Our prior findings suggest that ERGIC-localized Rab1 proteins can enhance the cargo-TMED10 complex, promoting TMED10 homo-oligomerization (Sun et al., 2024). Consistently, Rab1A and Rab1B, but not the UcPS-deficient mutants, expression enhanced TMED10 stability (Fig. 7, J-M), underscoring the critical role of endogenous cargo-TMED10 interactions enhanced by Rab1s in maintaining TMED10 homo-oligomer structure. These findings demonstrate that UcPS-competent cargo can stabilize TMED10 homo-oligomers even in the absence of other TMED family members.

Collectively, these findings support the existence of two distinct TMED oligomer populations that are differentially localized to the ER and ERGIC, participating in classical transport and UcPS, respectively. The transition from hetero-tetrameric to homo-oligomeric states is modulated by subcellular localization and interaction partners, with UcPS cargo interactions further stabilizing homo-oligomeric structures to promote the formation of UcPS translocators and facilitate secretion (Fig. 7 N).

## Discussion

Our research reveals that TMED proteins, both individually and in collaboration, play a pivotal role in governing the release of a diverse array of cargoes in UcPS. This regulatory capacity provides a means for fine-tuning UcPS in response to varying cellular contexts and physiological conditions. By employing secretome analysis, we have identified 710 potential UcPS cargoes, each of which exhibits distinct secretion regulation patterns mediated by different TMED proteins. Notably, the majority of these cargoes have been identified in human plasma, suggesting their physiological involvement in extracellular processes in humans (Fig. 1 E).

The extracellular functions of a substantial proportion of these secretory cargoes have not been previously reported, presenting an exciting opportunity to identify novel cytokines, hormones, or biomarkers. Prior studies have demonstrated that UcPS cargoes can function as cytokines, directly triggering signaling pathways, e.g. Interleukin 1s, histones, and FGF2 (Dinarello, 2018; Pallotta and Nickel, 2020; Silk et al., 2017). Indeed, our proteomic analysis has also uncovered multiple histones whose release is modulated by TMEDs (Fig. 1 H and Table S1). Additionally, UcPS factors can serve as extracellular modulators, participating in the remodeling of the extracellular environment (e.g., FABP4 and Galectins) (Popa et al., 2018; Prentice et al., 2021; Villeneuve et al., 2018) or being internalized by neighboring cells, thereby directly influencing intracellular processes through their intracellular activities (e.g., Tau and α-synuclein) (Luk et al., 2012; Reyes et al., 2013). The newly identified UcPS cargoes may fall into these established categories or operate through previously undiscovered mechanisms. Further investigations are warranted to clarify the extracellular functions of these cargoes.

TMED proteins possess several homologous domains, including the CT region, GOLD domain, and coiled-coil domain. The CT has been shown to bind COPI and COPII components, facilitating trafficking between the ER and Golgi, while the GOLD and coiled-coil domains regulate TMED oligomerization (Pastor-Cantizano et al., 2016). Notably, the CT also plays a critical role in binding cargoes, which is essential for initiating their translocation (Zhang et al., 2020). This study reveals that the CT regions of different TMEDs contribute to cargo specificity. Short CTs may function as ligand-like motifs for cargo binding, exhibiting specificity as point mutations in the cargo-binding surfaces, such as those in disease-associated PCBD1 mutants, disrupt interactions with TMED10, affecting cargo secretion.

Interestingly, secretome analysis shows overlapping cargo specificity among TMEDs, likely due to shared features of TMED-CTs, including conserved motifs for COPI and COPII binding. While the similarities and differences among TMED-CTs may explain both shared and distinct cargo recognition, the GOLD and coiled-coil domains may also play a role in cargo selection. Cargo interactions can influence TMED oligomerization, which is critical for translocation, and the GOLD and coiled-coil domains may impact the formation and structural conformation of TMED oligomers. These domains likely need to be precisely regulated to accommodate the specific translocation requirements of different cargoes.

Our findings demonstrate that the ERGIC is a pivotal site for TMED proteins to facilitate cargo translocation in UcPS. Although TMEDs continuously cycle between the ER, ERGIC, and Golgi (Dominguez et al., 1998; Gommel et al., 1999; Pastor-Cantizano et al., 2016), their translocation activity is specifically functional at the ERGIC. This specificity is likely because translocation at the ERGIC avoids both N- and O-glycosylation, which are initiated in the ER and Golgi. Glycosylation can potentially alter the properties of UcPS cargoes, rendering them inactive (Wegehingel et al., 2008). Therefore, translocation at the ERGIC maintains the functionality of these cargoes by preventing such modifications. Additionally, the ERGIC has been implicated in Golgi-bypass pathways, enabling direct trafficking from the ERGIC to endosomes or the plasma membrane (Saraste and Marie, 2018). This makes the ERGIC an optimal site for cargo translocation and subsequent Golgi-independent transport to the cell surface.

Systematic studies on UcPS using the yeast model system have identified Grh1 (the mammalian homologs of GRASP55 and GRASP65) as a central regulator of UcPS (Duran et al., 2010; Kinseth et al., 2007). A compartment of UcPS (CUPS) has been characterized, with tubulovesicular structures that localize near the ER exit sites (ERES) (Bruns et al., 2011; Cruz-Garcia et al., 2014). These structures share both structural and localization similarities with the UcPS-associated regions of the ERGIC. Notably, recent studies have shown that Grh1 co-localizes with ERGIC markers, suggesting that the Grh1-positive CUPS may represent an ERGIC-related compartment (Tojima et al., 2024). Furthermore, Grh1-positive CUPS are implicated in facilitating Golgi-bypass trafficking, potentially through membrane contacts with a modified pool of the trans-Golgi network (TGN) (Curwin et al., 2025). Thus, the THU pathway in mammalian cells may represent a homologous mechanism to yeast Grh1-mediated UcPS. A critical next step is to determine whether TMEDs act as translocators that facilitate cargo entry into the CUPS in yeast.

Importantly, our findings, in conjunction with previous and parallel researches, underscore the multifaceted regulation of TMED family proteins influenced by their subcellular location, lipid milieu, and interactions with partner molecules. This multifunctionality is evident in their roles related to GPI-anchored protein transport and UcPS cargo translocation. When situated within the ER, TMEDs predominantly assemble into stable tetramers, which engage with GPI-anchored proteins, facilitating their transit to the ERGIC and the Golgi apparatus. Conversely, while at the ERGIC, TMEDs have the propensity to form homo-oligomers, influenced by ERGIC-localized Rab1s (Sun et al., 2024). In this context, they function as cargo translocators, orchestrating the initial stage of vesicle-dependent UcPS, subsequently involving secretory autophagosomes, multi-vesicular bodies, and lysosomes (Rabouille, 2017; Zhang and Schekman, 2013; Zheng and Ge, 2022). These homo-oligomers on the ERGIC may not represent a stable form of TMEDs which may be degraded via proteasomal or lysosomal pathways (Hickey et al., 2023; Liu et al., 2008). However TMED homo-oligomers can be stabilized by the presence of UcPS cargoes, poised for translocation (Fig. 7 N). Therefore, cargo-facilitated stabilization of TMED homo-oligomers serves as a dedicated mechanism to prevent excessive translocator activity in the absence of cargoes, which could otherwise disrupt intracellular ion or protein homeostasis.

Multiple UcPS cargoes have been observed to employ both vesicle-dependent and independent pathways for their release (Dimou and Nickel, 2018; Sitia and Rubartelli, 2018; Zheng and Ge, 2022). For instance, IL-1β, which was utilized as a secretory cargo in our study, can be discharged via diverse mechanisms, including secretory autophagosomes, endo-lysosomes, exosomes, Gasdermin D-mediated pore formation on the plasma membrane, or direct plasma membrane translocation (Claude-Taupin et al., 2018; Evavold et al., 2018; Pallotta and Nickel, 2020; Qu et al., 2007; Shi et al., 2015; Sitia and Rubartelli, 2018). The predominant pathway for the release of a UcPS cargo is likely determined by specific physiological or pathological conditions, and further investigations are necessary to elucidate the factors governing these choices.

## Data and material availability

All data are available in the main text or the supplementary materials. Further information and requests for reagents should be directed to Lead Contact, Liang Ge. Plasmids and cell lines generated in this study will be made available upon request. We may require a payment and/or a completed Materials Transfer Agreement in case there is potential for commercial application.

## Acknowledgement

Dr. Liang Ge and Dr. Min Zhang are deeply grateful for the postdoc training done in Dr. Randy Schekman’s lab at University of California, Berkeley. We thank Dr. Bao-Liang Song (Wuhan University), Dr. Yongqiang Deng (South Medical University, China), Dr. Yong-Hui Zhang, Dr. Jing-Ren Zhang, Dr. Sen-Fang Sui and Dr. Shan Sun (Tsinghua University) for reagents. We thank Dr. Ye-Guang Chen and Dr. Li Yu (Tsinghua University) for helpful suggestions on the study. We thank Yimin Zheng (Austrian Academy of Sciences), Siqi Xu and Kun Zhu for technique assistance.

## Author contribution

Conceptualization: MZ, LG; Methodology: JZ, HW, YS, PC, XD, LZ , LZ, KZ, HD, MZ, LG; Investigation: JZ, HW, MZ, LG; Funding acquisition: MZ, LG; Project administration: MZ, LG; Supervision: MZ, LG; Writing – original draft: LG; Writing – review & editing: JZ, HW, MZ, LG.

## Supplemental information

Table S1. Mass spectrometry data for the secretome analysis

## Material and Methods

### Cell culture and transfection

HEK293T, U2OS, and HeLa cells were maintained in Dulbecco’s modified Eagle’s medium (DMEM) supplemented with 10% FBS at 37°C in 5% CO2. For immunofluorescence, cells were grown on coverslips (CITOGLAS). Transfection of plasmids into cells was performed using PEI (Polysciences, Inc.) for HEK293T and NEOFECT (Neofect (Beijing) biotech Co. Ltd.) or X-tremeGENE HP (Roche) or Sage LipoPlus (Sage (Beijing) Chemical Co. Ltd.) for U2OS and HeLa according to the manufacture’s protocols.

### Plasmids

The TMEDs family members (TMED7, TMED9, and TMED10 were purchased from DNASU, and the others were amplified from HEK293T cDNA) were PCR amplified and inserted into the FUGW vector with a V5 tag at the C-terminus. The TMEDs truncations and mutations were generated by mutagenesis PCR. The TMEDs fragments were PCR amplified and inserted into the pGEX-4T-1 vector. EGFP-tagged ERGIC-53, Streptavidin, and Streptavidin binding peptide plasmids were described previously (Li et al., 2022). LMAN2, HSP90AB1, HSP90B1, mIL-1β, FGF2, mIL-1α, mIL-33, mIL-36α, mIL-37, HSPB5, Galectin-1, Galectin-3,

Annexin A1, mIL-36RA, Tau and Rab1s plasmids were described previously (Sun et al., 2024; Zhang et al., 2020). RANGRF, PCBD1, NUDT14, HINT3, RAB3IL1, CA8, ELMO1 (isoform 2), CDKN1B, CNN1, BRMS1 were PCR amplified from cDNA and inserted into FUGW vectors for mammalian cell expression.

### Reagents and antibodies

We obtained QUANTI-Blue from InvivoGen, proteinase K, protease inhibitor cocktail, biotin, Duolink PLA kit, rabbit anti-ERGIC53, mouse anti-V5 and mouse anti-FLAG antibody from Sigma, phosphatase inhibitors cocktail from Roche, Phenylmethylsulfonyl fluoride (PMSF) from Amresco, Optiprep from Axis-Shield, and PNGaseF from NEB. We purchased Ni NTA Beads 6FF and Glutathione Beads 4FF from Smart-Lifesciences. We purchased mouse anti-HSP90AB1 antibody from Santa Cruz, rabbit anti-GRP94, anti-TMED2, anti-TMED3, anti-TMED4, anti-TMED9 and anti-TMED10 antibody from Proteintech, rabbit anti-TMED1, anti-TMED5, anti-TMED6, anti-TMED7 antibody from Novus Biologicals, cycloheximide (CHX), rabbit anti-V5, anti-HA and anti-GST antibody from CST, mouse anti-GM130 antibody from BD Biosciences, mouse anti-tubulin and anti-actin antibody from Abcam. Rabbit anti-RPN1 and anti-GFP antibody was described previously (Li et al., 2022; Ma et al., 2022; Zhang et al., 2020).

### Determination of cargo secretion

For secretion determination, cell culture medium was replaced with DMEM (TMED10-KO HEK293T, U2OS) or EBSS (wild-type HEK293T) containing indicated drugs for 1-2 hours. The medium was collected and concentrated (20-fold) by a 10 kDa Amicon filter (Millipore). Cells were lysed in SDS-PAGE loading buffer. Immunoblot was performed to determine the amounts of cargoes in the medium and cell lysate. SEAP activity (InvivoGen) was determined according to the manufacturers’ protocol.

### Lentiviral transduction and generation of CRISPR KO cells

For lentiviral transduction, FUGW plasmids containing the LMAN2/TMEDs-V5, was transfected into HEK293T cells along with lentiviral packaging plasmids pMD2.G and psPAX2 (Addgene) using PEI to produce lentiviral particles to infect U2OS cells.

For the generation of TMED10-KO cell lines in HeLa, the cells were transfected with pX330 containing TMED10 targeting sequences (sgRNA sequences: TCCGGCGCGGTTGAGGCCTT & TAACGGAAAAGGGCCGCGCC).

### Secretome analysis

The method for preparing conditioned medium was reported previously with minor modifications (Capaci et al., 2020; Georgilis et al., 2018). U2OS cells stably expressing LMAN2/TMEDs-V5 (TMED1-5 & 7, 9, 10-V5) were seeded in 15 cm dishes. 2 days later, the medium was diluted to 5% FBS with DMEM to reduce protein contamination in serum. The following day, cells were washed three times with PBS, and replaced with DMEM supplemented with 0.5 μg/ml Brefeldin A, an inhibitor of classical secretion. 2.5 hours later, the medium was collected, centrifuged at 2000 × *g* for 10 min to remove cellular debris, and filtered through a syringe filter (Pall Corporation). The clarified medium was then concentrated about 300-fold using 10 kDa filter tubes (Millipore) and the buffer was exchanged to 8M urea dissolved in PBS through three repeated dilutions and centrifugations. The protein concentration of the samples was determined using the BCA Protein Assay Kit (Beyotime), and an equivalent amount of each sample (100 μg) was proceeded to tandem mass tag (TMT)-based quantitative mass spectrometry in Protein Research and Technology Center at Tsinghua University.

### Membrane flotation assay

Membrane flotation assay was performed based on a previous study with certain modifications (Zhang et al., 2015). TMED10-KO HEK293T cells (100-mm culture dish) were transfected with TMEDs (TMED1-7 & TMED9, 10) and cargoes (mIL-1β-FLAG, galectin-3). After 24 h, cells were harvested and washed with ice-cold PBS. Following a centrifugation step at 600 × g for 5 minutes, the cell pellets were resuspended in HB1 lysis buffer (20 mM HEPES-KOH, pH=7.2, 400 mM sucrose, 1 mM EDTA, protease and phosphatase inhibitors, 0.3 mM DTT) and then lysed by passing through a 22G needle until around 85% lysis analyzed by Trypan Blue staining. The homogenates were centrifuged at 3,000 × g for 10 min and then the supernatant was ultra-centrifuged at 100,000 × g for 30 min. The membrane pellet was resuspended in 100 μL of B88 buffer (20 mM HEPES-KOH, pH=7.2, 250 mM sorbitol, 150 mM potassium acetate, 5 mM magnesium acetate, protease and phosphatase inhibitors, 0.3 mM DTT) combined with 200 μL of OptiPrep to achieve a final concentration of 40%. This was overlaid with 600 μL of 30% OptiPrep (diluted in B88 buffer) and 100 μL of B88 buffer, and then centrifuged at 45,000 rpm for 2 hours. The membrane fraction floating on the top (150 μL) was collected and subjected to immunoblot analysis.

### Cell-free translocation

The cytosol of wild-type HEK293T cells was prepared as previously described (Ge et al., 2013; Zhang et al., 2015). For membrane separation, the cells were transfected with plasmids encoding different TMEDs (TMED1-7 & 9, 10) or variants of TMEDs using PEI (polyethylenimine). 24 hours later, the cells were harvested and washed once with ice-cold PBS, and then lysed in HB1 lysis buffer by passing through a 22G needle. The lysate was centrifuged at 3,000 × g for 10 min and the supernatant was ultra-centrifuged at 100,000 × g for 30 min to collect the membrane pellet. The pellet was washed with B88 buffer for buffer exchange to B88, and then resuspend with B88 lysis buffer containing the cytosol of wild-type HEK293T cells (4 mg/ml final concentration). The phosphatidylcholine (PC) concentration was measured using a microplate spectrophotometer and adjusted to an equal concentration.

For the cell-free mIL-1β, RANGRF, and PCBD1 translocation, recombinant proteins (10 μg T7-mIL-1β-FLAG protein, 10 μg RANGRF-FLAG protein, or 30 μg PCBD1-FLAG protein in 120 μL reaction system), ATP regeneration system (40 mM creatine phosphate, 0.2 mg/ml creatine phosphokinase, and 1 mM ATP final) and GTP (0.15 mM final) were added to the purified membrane solution containing 4 mg/ml cytosol, then incubated at 30°C for 1.5 h. After then, the 100 μL reaction system was combined with 200 μL 60% OptiPrep to adjust the final concentration of Optiprep to 40%, followed by overlaying with 600 μL 30% OptiPrep (diluted in B88 buffer), 100 μL B88 buffer, and centrifuged at 45,000 rpm for 2 h. The membrane fraction floating on the top (150 μL) was collected and aliquoted into three fractions. The first fraction was a control, the second and third fractions were digested by protease K (20 μg/ml) without or with 1% Triton X-100 for 20 min on ice, with a total volume of 50 μL for each reaction. The reactions were stopped by 2 mM PMSF and incubated for 10 min on ice. Then 2 × SDS loading buffer was added and the samples were heated at 100°C for 10 min followed by immunoblot analysis.

### Immunofluorescence

Immunofluorescence was described previously (Zhang et al., 2020). Briefly, cells were washed twice with PBS and then incubated with 4% paraformaldehyde at room temperature for 20 min. The cells were washed twice with PBS and permeabilized with 0.2% TritonX-100 diluted in PBS for 3 min. After washing with PBS 3 times, cells were blocked by 10% FBS diluted in PBS at room temperature for 1 h and followed by primary antibody incubation at room temperature for 2 h. Cells were washed with PBS 3 times and followed by secondary antibody incubation at room temperature for 1 h. Fluorescence images were acquired using the Olympus FV3000 confocal microscope or Nikon AX confocal microscope. Quantification of the images was performed using ImageJ software.

### Retention Using Selective Hooks (RUSH) system

TMED10-KO HEK293T or HeLa cells were co-transfected with the hooks (ER hook: streptavidin-KDEL, ERGIC hook: streptavidin-TMED10ΔGD-HA or streptavidin-LMAN2-HA) and reporters (SBP-TMEDs-V5). To release the reporters from the ER or ERGIC, biotin (Sigma, B4501) was added to the cultured cells at the indicated concentration and time points. Images (TMED10-KO HeLa) were acquired using the Olympus FV3000 confocal microscope. Cargo secretion and cell-free translocation were conducted with TMED10-KO HEK293T cells following the procedures described above.

### Protein purification

For the expression of GST-tagged TMED family proteins, cargo proteins (mIL-1β-FLAG, RANGRF-FLAG, PCBD1-FLAG), chaperone protein HSP90AB1, HSP90B1 (GRP94), the constructs were transformed into the *E.coli* strain BL21 (DE3) plysS. After transformation, a single colony was picked and grown in 600 μL LB medium at 37°C with shaking at 220 rpm for approximately 6 hours. The pre-culture was further expanded to 50 ml and incubated overnight under the same conditions. The next day this culture was scaled up to 1 L LB medium followed by incubating at 37°C with shaking at 220 rpm until the OD_600_ reached 0.6-0.8. 100 μM IPTG was added to the culture medium and the protein expression was induced at 16°C with shaking at 160 rpm overnight. After expression, the bacterial pellets were collected and stored at -80°C for subsequent purification.

For purification of GST-tagged TMED family proteins, cell pellets were lysed in lysis buffer (50 mM Tris 8.0, 5 mM EDTA,150 mM NaCl,10% glycerol, 0.5 mg/mL lysozyme, 0.3 mM DTT and 1 mM PMSF) and then rotated at 4°C for 0.5 h. TritonX-100 was added to adjust to 1% final concentration and incubated for an additional 10 minutes. The lysates were sonicated and then centrifuged at 20,000 rpm for 1 h. The supernatant was collected and incubated with 0.5 mL of Glutathione Sepharose 4FF beads (Cytiva) at 4°C for 2 h. After incubation, the beads were washed with 15 volumes of wash butter (PBS with 0.1% TritonX-100). The proteins were eluted with elution buffer (50 mM Tris 8.0, 250 mM KCl, 25 mM glutathione, and 0.1% TritonX-100). The proteins were concentrated by Amicon® Ultra Filters (Merck) and further purified by size-exclusion chromatography (SEC) on a Superose 6 increase column (Cytiva) in SEC buffer containing PBS with 0.1% TritonX-100. The proteins were snap-frozen by liquid nitrogen and stored at - 80°C. For purification of HSP90AB1, all procedures were carried out without the addition of 0.1% Triton X-100.

For purification of his-tagged cargo protein mIL-1β, RANGRF, PCBD1, and chaperone protein HSP90B1 , cell pellets were lysed in lysis buffer (2x PBS, 10 mM imidazole, 0.5 mg/mL lysozyme, 0.3 mM DTT and 1 mM PMSF) and then rotated at 4°C for 0.5 h. The lysates were sonicated and then centrifuged at 20,000 rpm for 1 h. The supernatant was taken out and added 0.5 mL Ni-NTA agarose to incubate at 4°C for 2 h. The agarose was washed with 10 volumes of wash buffer containing 0.1% Tween20 and wash buffer (2 x PBS, 25 mM imidazole) followed. The proteins were eluted with elution buffer (2 x PBS, 250 mM imidazole). The proteins were concentrated by Amicon® Ultra Filters (Merck) and desalted by PD MiniTrap G-25 (Cytiva). The proteins were snap-frozen by liquid nitrogen and stored at -80°C.

For GST tag removal, 5 mL of the eluted proteins were mixed with 10 units of thrombin and incubated with rotation at room temperature (25°C) overnight. The proteins were concentrated using Amicon® Ultra Filters (Merck) and desalted using PD MiniTrap G-25 (Cytiva). Finally, the proteins were snap-frozen by liquid nitrogen and stored at -80°C.

### Lipid extraction

For in vitro translocation assay, total lipids were extracted from HEK293T cells as follows. 50 dishes (100-mm culture dish) of HEK293T cells were collected and washed with PBS. After centrifuging 600 × g for 5 min, the cell pellets were resuspended in PBS (cell pellets: PBS =1: 1). 1.2 ml chloroform/methanol solution (chloroform: methanol=1: 2) was added to each 300 μL cell suspension and vortexed for 30 s. The mixtures were shaken at 180 rpm for 1 h at 37°C. The chloroform phase was collected and dried under a stream of nitrogen gas and then the lipids were further dried at 37°C for 1 h. Dried lipids were resuspended in HEPES-KAc buffer (20 mM HEPES (pH 7.2), 150 mM potassium acetate) and incubated at 220 rpm for 1 h at 37°C to resuspend thoroughly. The phosphatidylcholine content of the lipid solution was measured by a standard kit (Wako) to normalize lipid concentration. The lipids were aliquoted and stored at -80°C.

### In vitro translocation and protease K protection assay

Proteoliposome constitution was performed according to a previous study with some modifications (Zhang et al., 2020). 1 mg lipids for each reaction tube were frozen in liquid nitrogen and thawed in a 42°C water bath alternately 10 times. Triton X-100 was added to achieve a final concentration of 0.05%, and the mixture was rotated at 4°C for approximately 30 minutes. Next, 10 μg recombinant proteins were added to the lipid solution (400 μL solution contain 10 μg recombinant proteins, 1 mg lipids, and 0.05% TritonX-100) and the mixture was rotated at 4°C for 1 h. Each solution was incubated with 6-8 mg Bio-Beads SM2 Resin (Bio-Rad) which was pre-equilibrium by HEPES-KAc reaction buffer to remove the detergent and rotated at 4°C around 1 h. This step was repeated 5 times (10 mg beads used during the third incubation, which was extended overnight.). The Bio-Beads were removed at 1,500 × g centrifugation at 4°C each step and after the last time, the liposome solution froze in liquid nitrogen and thawed in a 42°C water bath alternately 5 times. To remove the free proteins, a membrane flotation step was performed. For each 350 μL solution, 350 μL 50% OptiPrep (diluted in HEPES-KAc buffer) was added. The mixture was overlaid with 560 μL 20% OptiPrep and 105 μL HEPES-KAc buffer, centrifuged at 100,000 × g for 2 h and the top 200 μL fraction was collected and diluted with 100 μL HEPES-KAc buffer as proteoliposome solution.

For the in vitro protein translocation, 10 μg cargo proteins (e.g. mIL-1β-FLAG, mIL-33-FLAG, mIL-33-K-FLAG) were added to proteoliposome solution (150 μL system) and then incubated at 30°C for 1 h. After incubation, 5 μL reaction buffer was taken as input and the left was added 150 μL 50% OptiPrep (diluted in HEPES-KAc buffer). The mixture was overlaid with 240 μL 20% OptiPrep and 45 μL HEPES-KAc buffer, centrifuged at 45,000 rpm for 2 h and the 90 μL top fraction was collected and aliquoted into 3 fractions. All fractions were loaded to 50 μL. The first fraction served as a control, the second and the third fractions were digested by 10 μg/ml protease K without or with 0.1% TritonX-100 for 20 min on ice. The reactions were stopped by 1mM PMSF and incubated for 10 min on ice. Finally, 2 × SDS loading buffer was added and the samples were heated at 100°C for 10 min followed by immunoblot analysis.

For the temperature tests, translocation systems were incubated at either 0°C or 30°C for 1 h. For the protein unfolding assay, cargo proteins were pre-treated with an equal volume of 8 M urea for at least 30 min. For chaperone-associated translocation, HSP90B1 was added with recombinant proteins simultaneously, HSP90AB1 and ATP (1.5 mM final concentration) were added with cargo proteins before translocation.

### In situ Duolink

Duolink PLA kit was purchased from sigma and the assay was performed according to the product manual. To investigate interactions between TMEDs and cargoes, variants of TMEDs and cargoes were co-transfected into HeLa cells. 24 hours post transfection, the cells were washed twice with PBS and then incubated with 4% paraformaldehyde at room temperature for 20 min. The cells were washed twice with PBS and permeabilized with 0.2% TritonX-100 diluted in PBS for 3 min. The cells were blocked, incubated with primary antibodies and PLA probes followed by ligation and amplification using the recommended conditions according to the manual. Images were acquired using the Olympus FV3000 confocal microscope and Quantification was performed using ImageJ.

### LC-MS/MS analysis

For the label-free MS quantification of TMEDs, cellular membrane fractionation was performed to enrich the TMEDs. Briefly, U2OS or HeLa cells were harvested and washed twice with ice-cold PBS, and then lysed in HB1 lysis buffer by passing through a 22G needle. The lysate was centrifuged at 1,000 × g for 10 min and the supernatant was ultra-centrifuged at 100,000 × g for 30 min to collect the whole membrane pellet. The pellet was washed twice with PBS before resuspended with PBS. Then the suspension was digested with PNGaseF for 2.5 h to remove the N-glycans in certain TMEDs (TMED1, 4, 7, 9) to avoid the interference with MS quantification. After the immunoblot verification that all N-glycans in TMEDs (TMED1, 4, 7, 9) were removed, the remaining samples were proceeded to SDS-PAGE and the gel between 15-35 kDa containing all TMEDs was collected for MS detection.

### Gel Filtration Chromatography

Gel filtration assay was performed based on a previous study with certain modifications (Ma et al., 2022; Marzioch et al., 1999). One dish (100-mm culture dish) of WT HeLa cell was harvested and washed with PBS 3 times. Following a centrifugation step at 600 × g for 5 minutes, the cell pellets were resuspended and lysed in 700 μL of lysis buffer (4% octyl glucoside, 100 mM NaCl, 10% glycerol, 1 mM PMSF, and 20 mM HEPES, pH 7.4) at 4°C for 1 h. The cell supernatant was then isolated after centrifugation at maximum speed at 4°C for 15 min. 500 μL of the supernatant was applied to Superdex 200 Increase 10/300 GL gel filtration column and each 500 μL fractions were collected. This prosses was performed in a buffer containing 1% octyl glucoside, 100 mM NaCl, 10% glycerol, 1 mM PMSF, and 20 mM HEPES, pH 7.4. The collected fractions were calibrated by western blot using antibodies against endogenous TMED protein.

### Co-immunoprecipitation

For the Co-IP using L and P fraction, the membrane fractionation protocol was reported previously (Ge et al., 2013; Li et al., 2022). Briefly, the cells were harvested and lysed in HB1 lysis buffer. The homogenate was centrifuged at 3,000 × g for 10 min and the supernatant was ultra-centrifuged at 25,000 × g for 20 min to collect the 25k pellet. The 25k membrane was resuspended with 0.45 ml 1.25 M sucrose buffer overlayed with 0.3 ml 1.1 M and 0.3 ml 0.25 M sucrose buffer (Golgi isolation kit; Sigma) and then centrifuged at 120,000 × g for 2 h (TLS 55 rotor, Beckman). Following this, two fractions were collected: one at the interface between 0.25 M and 1.1 M sucrose (L fraction), and the other from the pellet at the bottom of the tube (P fraction). These fractions were lysed in Co-IP buffer (50 mM Tris, 150 mM NaCl, 1 mM EDTA, 0.5% NP40, pH 7.2) with protease inhibitors and DTT, and incubated with anti-V5 agarose at 4°C for 3 h. Then the agaroses were washed four times with Co-IP buffer followed by immunoblot.

### CHX chase assay

Based on our previous study (Ma et al., 2022), TMED10-KO HEK293T or HeLa cells were transfected with indicated plasmids. 24 hours later, cells were treated without or with 50 µg/ml CHX as indicated and were collected at each indicated time point for immunoblot analysis.

### Quantification and statistics

For the quantification of cargoes secretion, the relative level of cargoes in medium was normalized to cargoes level in corresponding cell lysate. For the quantification of in vitro translocation, the relative cargoes translocation efficiency was normalized to TMEDs level. The statistical information of each experiment, including the statistical methods, the P-values, and numbers (n), were shown in the figures and corresponding legends. Statistical analyses were performed in GraphPad Prism.

## Data and code availability

This study did not generate any unique datasets or code.

## Additional resources

This study did not generate any additional resources.

---

**Figure S1.**
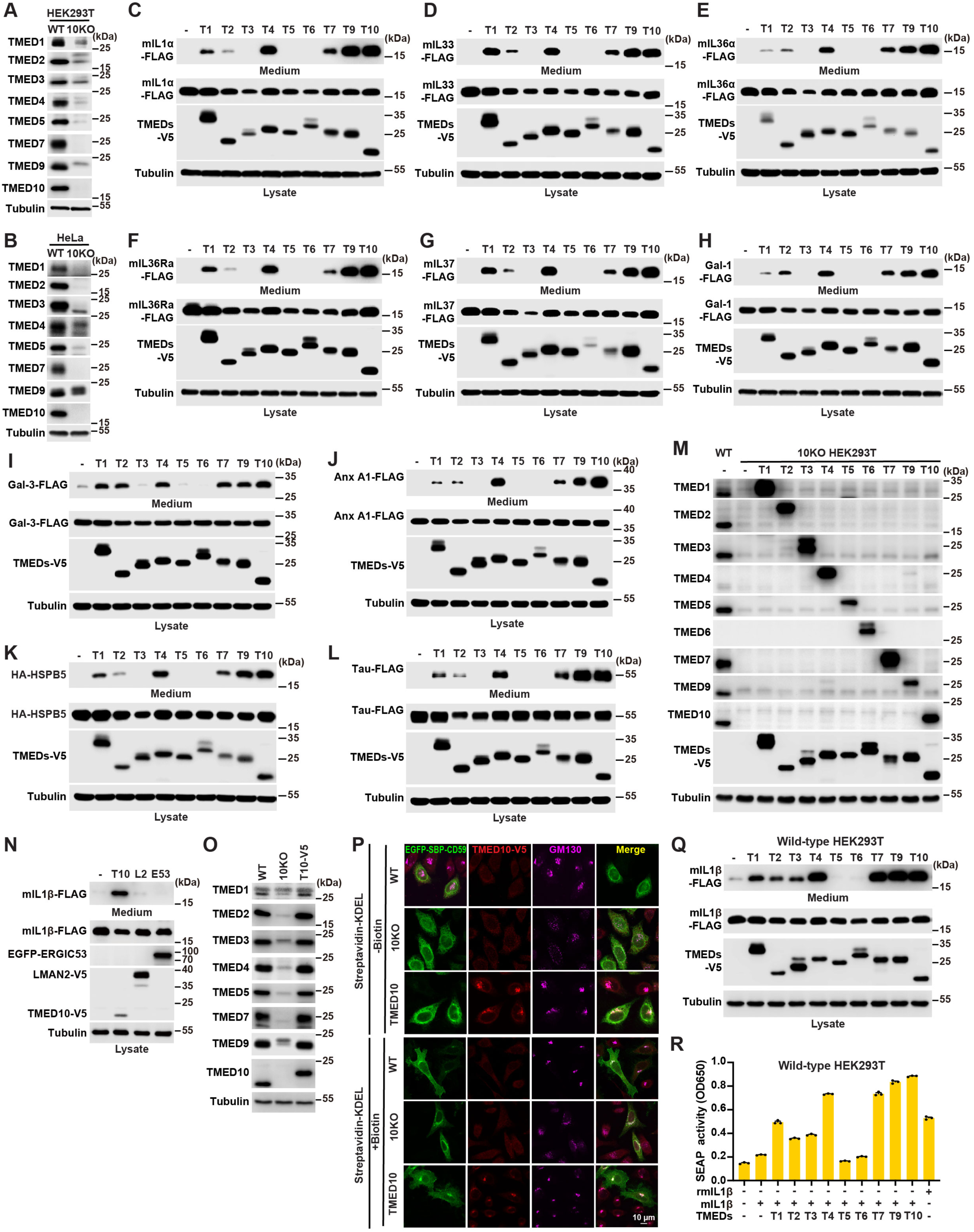
Anaylsis of TMEDs function in UcPS and CD59 trafficking. **(A and B)** Expression level of endogenous TMEDs (TMED1-5 & 7, 9, 10) in wild-type (WT) or TMED10 knockout (10KO) HEK293T (A) and WT or 10KO HeLa cell lines (B) generated by CRISPR-Cas9. The endogenous TMED6 level was low and undetectable in these cell lines. The data are representative of three independent experiments. **(C-L)** Secretion of mIL-1α (C), mIL-33 (D), mIL-36α (E), mIL-36Ra (F), mIL-37 (G), Galectin-1 (H), Galectin-3 (I), Annexin A1 (J), HSPB5 (K) and Tau (L) in TMED10-KO HEK293T cells with control or expression of TMEDs (TMED1-7 & 9, 10). The data are representative of three independent experiments. **(M)** Level of endogenous and exogenous TMEDs in WT or TMED10-KO HEK293T cells with control or TMEDs (TMED1-7 & 9, 10) expression for 24h. **(N)** Secretion of mIL-1β in WT HEK293T cells with control or TMED10 (T10), LMAN2 (L2), ERGIC53 (E53) expression. The data are representative of three independent experiments. **(O)** Level of endogenous TMEDs (TMED1-5 & 7, 9, 10) in WT or TMED10-KO HEK293T cells with control or TMED10-V5 expression for 96h. **(P)** Surface delivery of CD59 analyzed by hook and release assay. WT and TMED10-KO HeLa cells with control or TMED10-V5 expression for 72 h were co-transfected with EGFP-SBP-CD59 and streptavidin-KDEL. 23 hours later, the cells were treated without or with 40 μM biotin for 1 h, and then immunofluorescence was conducted with anti-V5 and anti-GM130 antibody. Scale bar, 10 μm. **(Q)** Secretion of mIL-1β in WT HEK293T cells with control or TMEDs (TMED1-7 & 9, 10) expression. The data are representative of three independent experiments. **(R)** HEK-Blue IL-1β cells (InvivoGen) were treated with culture medium derived from indicated HEK293T cells or 0.1 μg/ml recombinant mIL-1β (rmIL-1β) as a positive control. Levels of SEAP indicating IL-1β activity in the medium were monitored using QUANTI-Blue (n=3).

**Figure S2.**
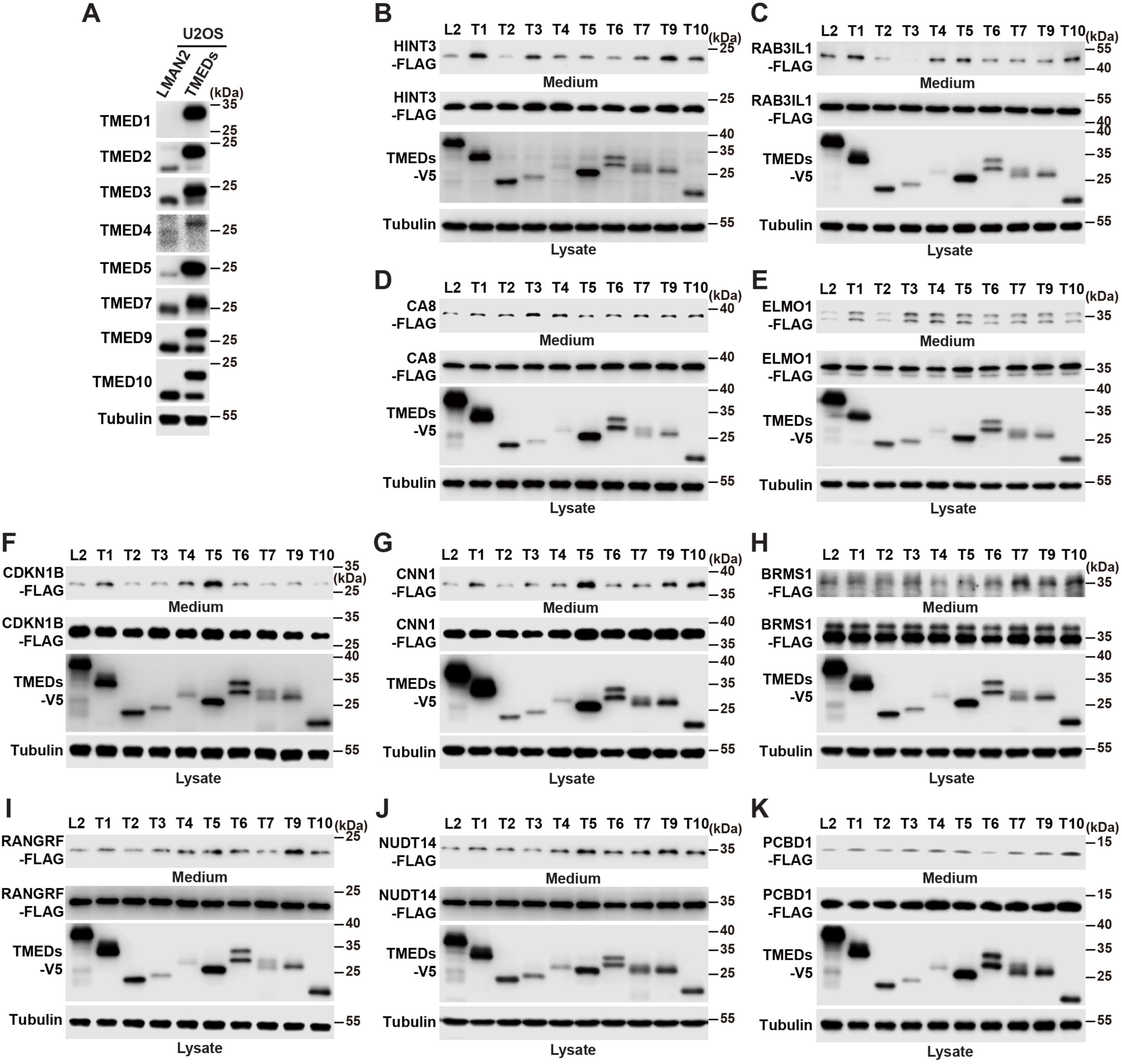
Confirmation of TMED-mediated selective cargo release. **(A)** Expression level of TMEDs (TMED1-5 & 7, 9, 10) in U2OS cells with stably LMAN2-V5 or TMEDs-V5 expression. **(B-K)** Secretion of HINT3 (B), RAB3IL1 (C), CA8 (D), ELMO1 (E), CDKN1B (F), CNN1 (G), BRMS1 (H), RANGRF (I), NUDT14 (J) , and PCBD1 (K) in U2OS cells with LMAN2-V5 or TMEDs-V5 (TMED1-7 & 9, 10) stable expression. The data are representative of three independent experiments.

**Figure S3.**
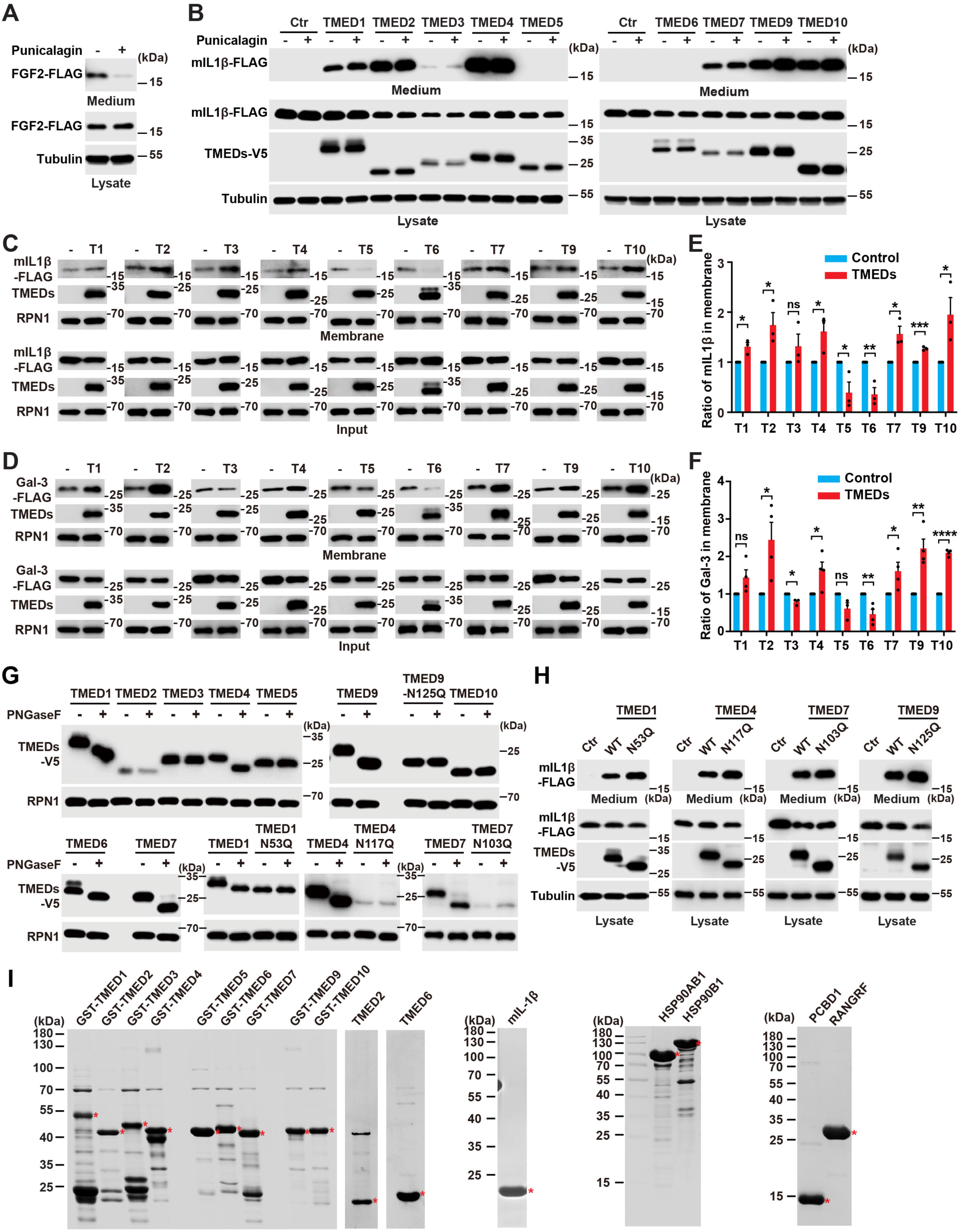
TMEDs mediate UcPS via membrane trafficking, TMEDs glycosylation and protein purification. **(A)** Secretion of FGF2 in TMED10-KO HEK293T cells without or with 10 μM punicalagin treatment for 1.5 h. The data are representative of three independent experiments. **(B)** Secretion of mIL-1β in TMED10-KO HEK293T cells with control or TMEDs (TMED1-7 & 9, 10) expression, without or with 10 μM punicalagin treatment for 1.5 h. The data are representative of three independent experiments. **(C and D)** Membrane floatation assay for the amounts of mIL-1β (C) and galectin-3 (D) in the membrane fractions and total cell lysates in TMED10-KO HEK293T cells with control or TMEDs-V5 (TMED1-7 & 9, 10) expression. **(E and F)** Quantification of the ratio of mIL-1β (E) and galectin-3 (F) (mean ± SEM) in membrane fractions as shown in (C) and (D), the control group was set as 1. p values were calculated by two-tailed t test (n ≥ 3). ns, non-significant; *P < 0.05; **P < 0.01; ***P < 0.001; ****P < 0.0001. **(G)** Glycosylation detection in TMED10-KO HEK293T cells expressing TMEDs (TMED1-7 & 9, 10) or TMEDs glycosylation site mutants (TMED1-N53Q, TMED4-N117Q, TMED7-N103Q, and TMED9-N125Q) without or with PNGaseF digestion. The data are representative of three independent experiments. **(H)** Secretion of mIL-1β in TMED10-KO HEK293T cells with control or expression of TMEDs-V5 (TMED1, 4, 7, 9) or its glycosylation site mutant (TMED1-N53Q, TMED4-N117Q, TMED7-N103Q, and TMED9-N125Q). The data are representative of three independent experiments. **(I)** Coomassie blue staining of GST-TMEDs (TMED1-7 & 9, 10), HSP90AB1, HSP90B1, mIL-1β-FLAG, PCBD1 and RANGRF expressed in E.coli expression system and GST-TMEDs (TMED2 & 6) after thrombin digestion to remove GST-tag.

**Figure S4.**
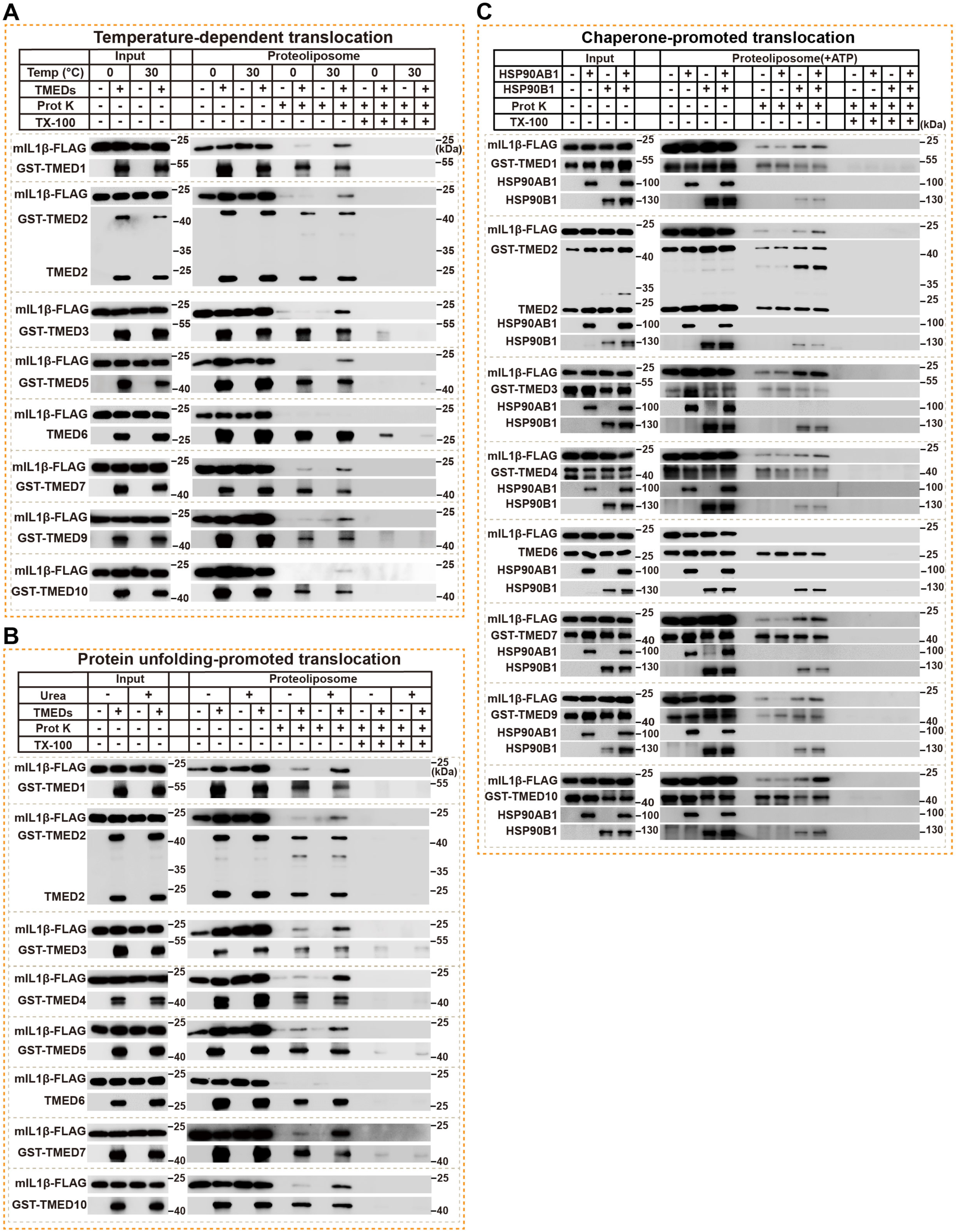
Accessory factors of TMEDs-mediated cargo translocation. **(A)** In vitro translocation assay with control or proteoliposomes composed of GST-TMEDs (TMED1, 3, 5, 7, 9, 10) or TMEDs (TMED2 & 6) performed at 0°C or 30°C. **(B)** In vitro translocation assay with control or proteoliposomes composed of GST-TMEDs (TMED1, 3, 4, 5, 7, 10) or TMEDs (TMED2 & 6) performed without or with 4 M urea pre-treatment to mIL-1β. **(C)** In vitro translocation assay with control or proteoliposomes composed of GST-TMEDs (TMED1, 3, 4, 7, 9, 10) or TMEDs (TMED2 & 6) performed in the presence or absence of HSP90s (HSP90AB1 outside and HSP90B1 inside) with ATP.

**Figure S5.**
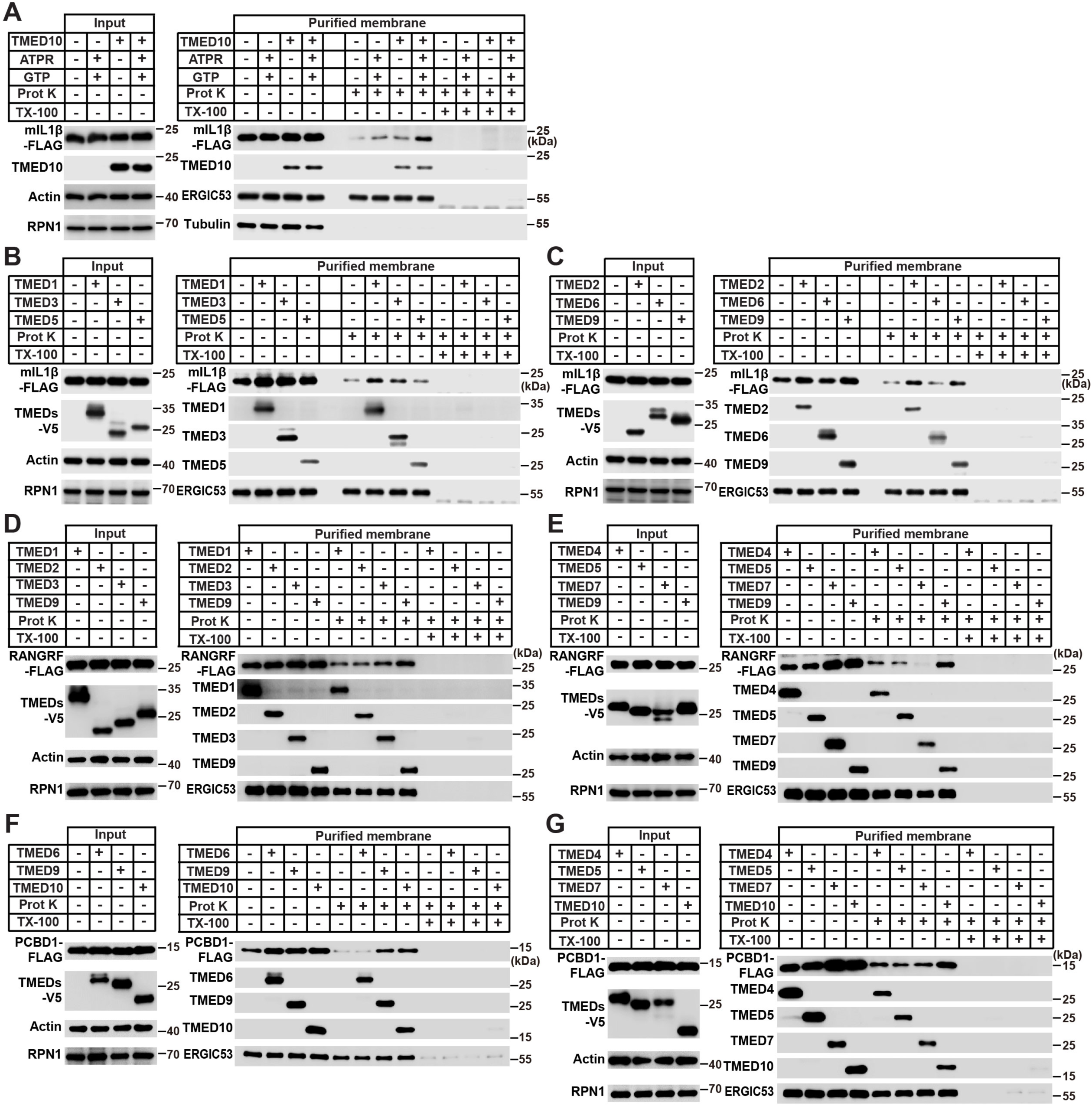
Cell-free translocation of cargoes by TMEDs. **(A)** Cell-free translocation of mIL-1β using membranes from TMED10-KO HEK293T cells without or with TMED10-V5 expression, without or with ATPR and GTP treatment. The data are representative of three independent experiments. **(B and C)** Cell-free translocation of mIL-1β using membranes from TMED10-KO HEK293T cells without or with TMEDs (TMED1, 3, 5 for (B), TMED2, 6, 9 for (C)) expression. **(D and E)** Cell-free translocation of RANGRF using membranes from TMED10-KO HEK293T cells with TMEDs (TMED1, 2, 3, 9 for (D), TMED4, 5, 7, 9 for (E)) expression. **(F and G)** Cell-free translocation of PCBD1 using membranes from TMED10-KO HEK293T cells without or with TMEDs (TMED6, 9, 10 for (F) TMED4, 5, 7, 10 for (G)) expression.

**Figure S6.**
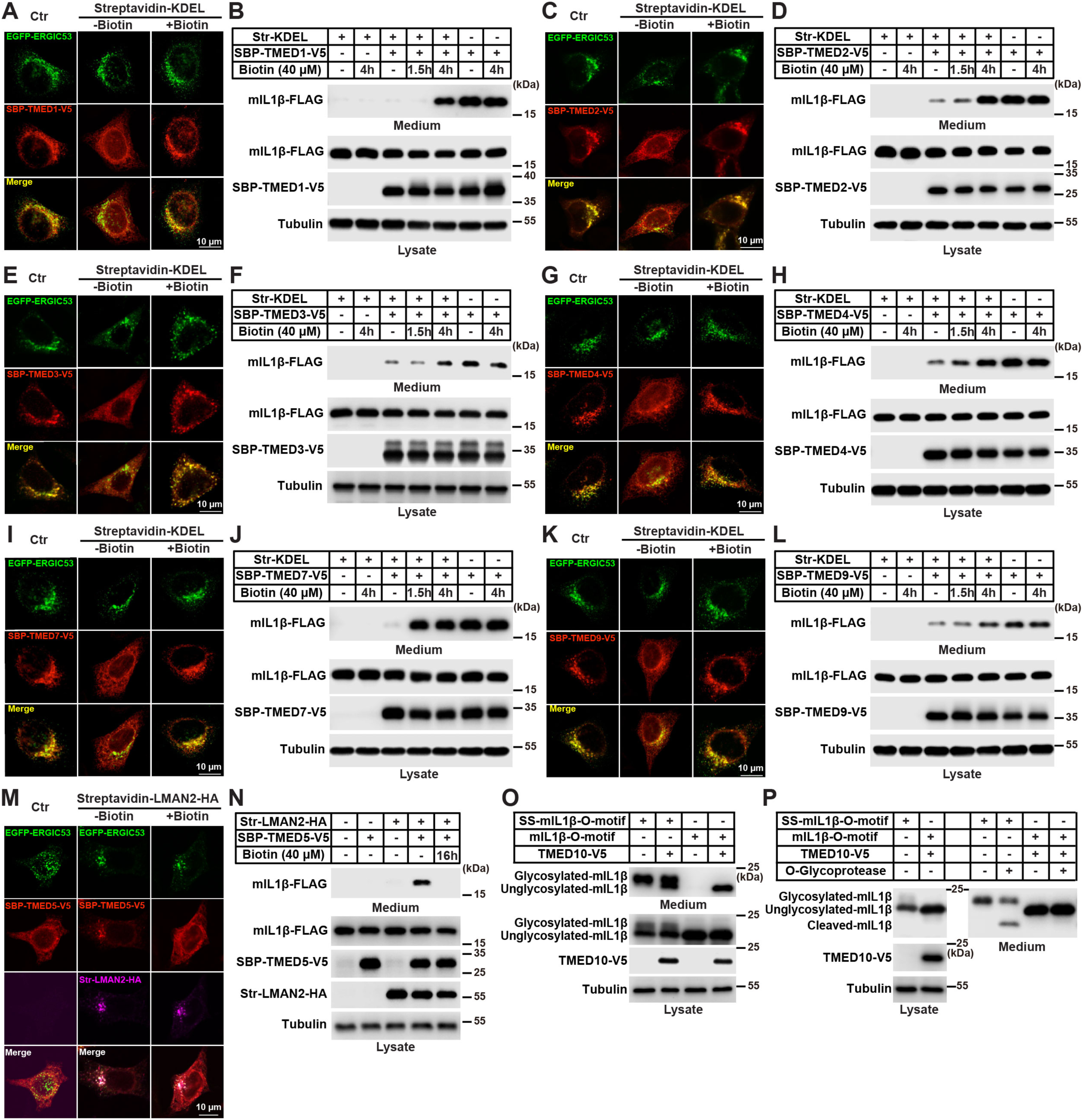
ERGIC localization is required for the function of TMEDs in UcPS. **(A, C, E, G, I and K)** TMED10-KO HeLa cells were co-expressed EGFP-ERGIC-53 and SBP-TMEDs-V5 (SBP-TMED1-4 & 7, 9-V5), without or with Str-KDEL expression, and treated without or with 40 μM biotin for 4 h. Immunofluorescence was performed with anti-V5 antibody. Scale bar, 10 μm. **(B, D, F, H, J and L)** Secretion assay of mIL-1β in TMED10-KO HEK293T cells combined with RUSH system. The cells were co-transfected with mIL-1β-FLAG, Str-KDEL, and SBP-TMEDs-V5 (SBP-TMED1-4 & 7, 9-V5) as indicated and treated without or with biotin at the indicated concentration and time points. The data are representative of three independent experiments. **(M)** TMED10-KO HeLa cells were co-expressed EGFP-ERGIC-53 and SBP-TMED5-V5, without or with Str-LMAN2-HA expression, and treated without or with 40 μM biotin for 16 h. Immunofluorescence was performed with anti-V5 and anti-HA antibody. Scale bar, 10 μm. **(N)** Secretion assay of mIL-1β in TMED10-KO HEK293T cells combined with RUSH system. The cells were co-transfected with mIL-1β-FLAG, Str-LMAN2-HA, and SBP-TMED5-V5 as indicated and treated without or with 40 μM biotin for 16 h. The data are representative of three independent experiments. **(O)** Secretion assay in HEK293T cells with control or TMED10-V5, and with conventional secretion cargo ss-mIL-1β-O-motif or UcPS cargo mIL-1β-O-motif expression (ss for signal peptide, O-motif for a reported O-GalNAc glycosylation motif with amino acid sequence GATGAGAGAGTTPGPG). The data are representative of three independent experiments. **(P)** Secretion and O-glycoprotease digestion assay in HEK293T cells with control or TMED10-V5, and with ss-mIL-1β-O-motif or mIL-1β-O-motif expression. The medium was collected and treated without or with O-glycoprotease IMPa (NEB). The data are representative of three independent experiments.

**Figure S7.**
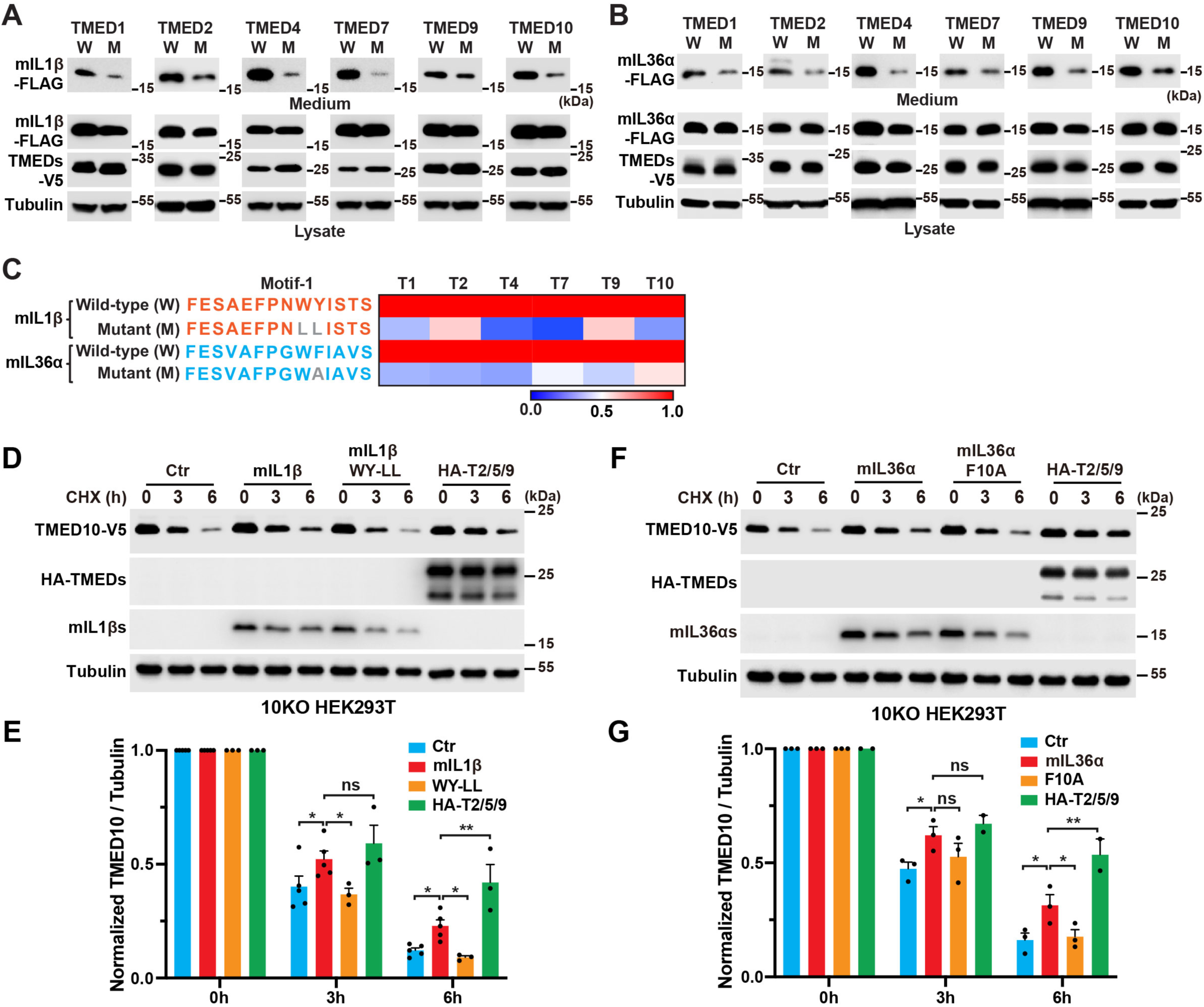
UcPS cargoes stablize the TMED10 homo-oligomer. **(A and B)** Secretion of wild-type mIL-1β (W) & mIL-1β-WY (9,10)-LL mutant (M), or wild-type mIL-36α (W) & mIL-36α-F10A (the site in the motif-1 identified previously) mutant (M) in TMED10-KO HEK293T cells with expression of TMEDs (TMED1, 2, 4, 7, 9, 10). The data are representative of three independent experiments. **(C)** Heatmap showing the relative secretion of mIL-1β & mIL-1β WY (9,10)-LL, and mIL36α & mIL36α-F10A in (A and B). **(D)** Turnover of TMED10-V5 in CHX chase assay without or with mIL-1β-FLAG, or its UcPS-deficient mutant (WY-LL) or three other subfamily members (HA-TMED2, 5, 9) co-expression in TMED10 KO HEK293T cells. **(E)** Quantification of normalized TMED10-V5 (mean ± SEM) as shown in (D), the relative level of TMED10-V5 was normalized to Tubulin and the 0 h control group was set as 1; p values were calculated by two-way ANOVA (n ≥ 3). **(F)** Turnover of TMED10-V5 in CHX chase assay without or with mIL-36α, mIL-36α-F10A and HA-TMED2, 5, 9 co-expression in TMED10 KO HEK293T cells. **(G)** Quantification of normalized TMED10-V5 (mean ± SEM) as shown in (F); p values were calculated by two-way ANOVA (n ≥ 2). ns, non-significant; *P < 0.05; **P < 0.01; ***P < 0.001; ****P < 0.0001.

